# Spatial Reorganization of Object Representations in High-Level Visual Cortex Distinguishes Working Memory from Perception

**DOI:** 10.1101/2025.06.29.662186

**Authors:** Wanru Li, Jia Yang, Pinglei Bao

## Abstract

The human visual system balances veridical object visual perception with flexible object visual working memory (VWM), both relying on high-level visual regions. However, how these competing demands shape spatial representations remains unclear. Here, we ask whether VWM inherits the spatial constraints observed in the lateral occipital complex (LOC) during perception, or instead reorganizes these representations to meet mnemonic demands. Using matched bilateral presentation paradigms and fMRI-based decoding, we systematically compared spatial representations during perception and VWM. This approach revealed a striking dissociation: during perception, object information is largely confined to the contralateral LOC, whereas during VWM, robust ipsilateral representations emerge—even when both hemifields must be remembered. Vertex-ablation analyses revealed that VWM engages 70–90% of ipsilateral LOC territories, far exceeding those recruited during unilateral perception. Neither increased attentional span nor top-down feedback from association areas fully explained this expansion; rather, ipsilateral LOC patterns closely mirrored contralateral sensory representations, implicating interhemispheric coordination in VWM. Together, these findings demonstrate that object VWM flexibly recruits distributed high-level visual cortex, with spatial reorganization distinguishing mnemonic flexibility from perceptual fidelity.

## Introduction

The human visual system faces two fundamental challenges: constructing accurate representations of the current environment through perception and maintaining behaviorally relevant information through visual working memory (VWM). Despite their distinct functional goals, perception and VWM recruit overlapping neural circuits (Yu & Shim, 2017; Emrich et al., 2013; Harrison & Tong, 2009; Serences et al., 2009). Specifically, activity in high-level visual cortex, for instance, the lateral occipital complex (LOC), supports object identity encoding during perception (Kriegeskorte et al., 2008; Schwarzlose et al., 2008; Haxby et al., 2001) and is also implicated in maintaining object representations during VWM (Xu, 2023; Lepsien & Nobre, 2007; Xu & Chun, 2006).

Although visual perception and VWM share neural substrates, they differ fundamentally in their computational demands. Visual perception emphasizes efficient sensory encoding of the external environment, typically through spatially constrained coding schemes. This is reflected in the retinotopic organization of early visual areas and contralateral biases observed in higher visual regions (Groen et al., 2022; Benson et al., 2018; Engel et al., 1997; DeYoe et al., 1996). In contrast, VWM must maintain internal representations without ongoing sensory input, often in the face of distraction or interference. This functional shift is thought to require more flexible or distributed coding strategies that prioritize stability over spatial specificity (Christophel et al., 2017; D’Esposito & Postle, 2015). These contrasting demands suggest that perception and VWM may impose different spatial constraints on shared cortical substrates, potentially leading to distinct patterns of representational organization.

Evidence from early visual cortex suggests that spatial coding is flexible and adapts to task demands. Perceptual representations in low-level visual areas adhere closely to retinotopic organization, with decodable sensory information confined to corresponding retinotopic regions (Tong et al., 2012). However, during VWM maintenance, this spatial lateralization effect becomes more flexible: contralateral preferences persist when memorizing bilateral stimuli (Pratte & Tong, 2014), but significantly diminish during unilateral memory maintenance (Zhao et al., 2021; Ester et al., 2009). These findings suggest that early visual cortex can shift from a strictly retinotopic code to a more unified mnemonic representation, depending on task context.

This raises an important question: Does similar spatial flexibility extend to higher-level regions like the LOC, which must balance spatial sensitivity with object-level abstraction? The LOC offers a unique opportunity to study spatial coding differences between perception and VWM, due to its hybrid spatial properties. On one hand, LOC exhibits spatial sensitivity: object representations vary with position, and robust contralateral biases have been consistently observed in fMRI studies (Silson et al., 2022; Reithler et al., 2017; Hong et al., 2016; Kravitz et al., 2010; Niemeier, 2004). On the other hand, LOC also exhibits position invariance, maintaining object identity representations even across visual hemifields (Ito et al., 1995). Unlike the strict retinotopic organization of EVC, this combination allows LOC to flexibly adjust to varying input configurations; for instance, bilateral object presentation results in near-absent ipsilateral representation compared to unilateral displays, consistent with dynamic competition for spatial processing resources (Bao & Tsao, 2018; Reithler et al., 2017). These findings suggest that during perception, LOC balances spatial selectivity and position invariance.

While these properties of the LOC are well established during perception, it remains unclear whether—and how—similar spatial coding principles operate during VWM. VWM requires maintaining internal representations without external input, which may rely on different coding strategies. Given the LOC’s dual sensitivity to spatial position and position-invariant identity, does it preserve the spatial constraints observed during perception, or does it adapt by recruiting additional ipsilateral resources? Moreover, is this flexibility stable across different levels of memory load?

To address these questions, we systematically compared spatial representational patterns in the LOC across perception and VWM using matched bilateral presentation paradigms. Participants viewed or memorized real objects across five conditions: unilateral perception, bilateral perception, unilateral VWM, bilateral VWM, and bilateral attention task. This design allowed us to examine how the representational structure—particularly the involvement of the ipsilateral hemisphere—varies across tasks.

Our results revealed a fundamental dissociation between perception and VWM in the spatial representation of objects within the LOC. Specifically, bilateral perceptual configurations result in negligible ipsilateral representation in the LOC, whereas VWM robustly engages ipsilateral regions even under high memory demands. This VWM-related expansion recruits additional cortical areas beyond those activated during unilateral perception or attentional tasks and adopts a sensory-like representational pattern. Although both perceptual and mnemonic representations share similar forms, these findings underscore spatial reorganization as a key factor distinguishing perceptual and mnemonic processing in high-level visual cortex.

## Results

To directly compare spatial coding in the LOC during perception and VWM, we designed a set of matched fMRI tasks involving unilateral and bilateral object presentation under perceptual, mnemonic, and attentional conditions. Using decoding, representational similarity analysis (RSA), vertex-ablation and searchlight RSA, we examined how the spatial structure of object representations in LOC flexibly adapts to different task demands. To determine whether these effects were specific to high-level visual areas, the early visual cortex (EVC) was included as a control region, given its well-characterized retinotopic organization. We also examined association areas, including the intraparietal sulcus (IPS) and prefrontal cortex (PFC), to investigate potential sources of sensory representations. Each participant completed three perception tasks, two VWM tasks, and one attention task under fMRI scanning for more than 11 hours. Regions of interest (ROIs), including the high-level object-selective LOC and EVC, were identified using independent localizer scans (see Figure S1b, c, and Methods).

### Absence of ipsilateral coding in the LOC during bilateral object perception

To establish baseline spatial representations in the LOC during perception, we compared responses to objects presented unilaterally (unilateral perception, UP; Figure 1a) versus bilaterally (bilateral perception, BP; Figure 1b). Twenty objects (Figure S1a) were used across all six tasks. In both UP and BP tasks, participants were required to maintain fixation by performing a fixation-change detection task. We then separately assessed contralateral and ipsilateral representations by decoding object identity using vertices from the ROIs within a single hemisphere. For contralateral representations, for example, we used right-hemisphere vertices to decode objects shown in the left visual field, and vice versa. A 20-class linear support vector machine (SVM) was trained for each decoding analysis, with decoding accuracy calculated via leave-one-trial-out cross-validation (see Methods).

**Figure 1.**
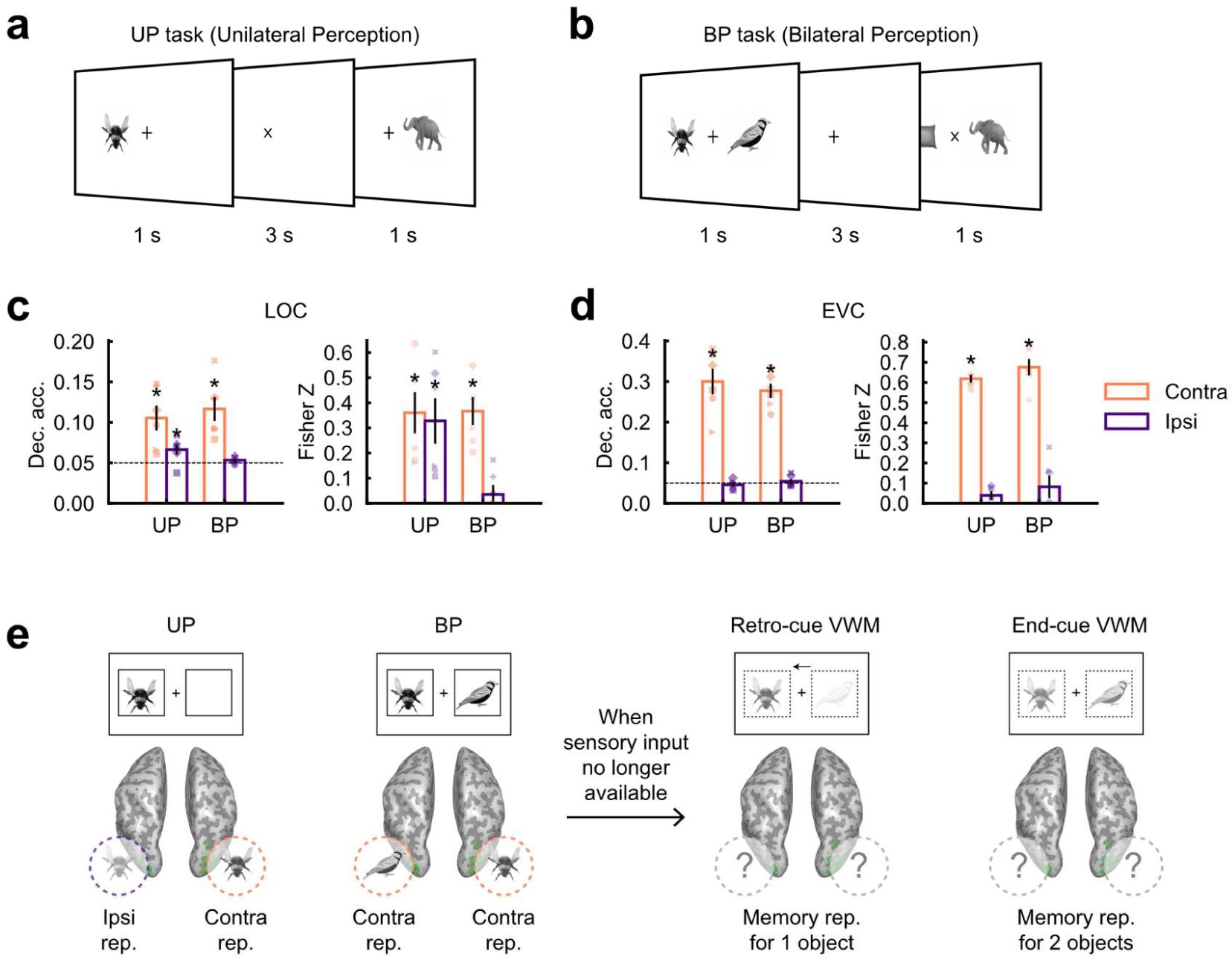
Perception task design and results. **a.** Illustration of the unilateral perception (UP) task. On each trial, a single object was presented on one side of the screen. Participants were required to maintain fixation and respond to any fixation change. **b.** Illustration of the bilateral perception (BP) task (Experiment 1b). Similar to the UP task, but two objects were presented simultaneously, one on each side of the screen. **c.** Decoding accuracy in UP / BP tasks (Left) and correlations between RDMs in UP / BP tasks and RDMs in CP task (Right) in LOC. Coral color represents the contralateral condition, while indigo color represents the ipsilateral condition. The horizontal dashed black lines indicate chance level. Error bars represent ± 1 *s.e.m.* across participants. Each symbol corresponds to an individual participant, with consistent symbol-to-participant mapping across all figures (see Figure S3). **d.** Same analysis as in **c.**, but for the EVC. **e.** Left, schematic depiction of LOC object representations during perception. The UP task yielded both contralateral and ipsilateral representations for object identity, whereas the BP task showed object representations only in the contralateral LOC. Right, an open question regarding the neural representation of objects during VWM when sensory input is no longer available, which was further examined in subsequent experiments under comparable encoding conditions. Contra, contralateral; Ipsi, ipsilateral; VWM, visual working memory; rep., representation; Dec. acc., decoding accuracy; *s.e.m.*, standard error of the mean. * *p* < 0.05, uncorrected, Wilcoxon signed-rank test.

In the UP task, decoding accuracy in the LOC significantly exceeded chance level for both contralateral and ipsilateral conditions (Figure 1c left, contralateral: *p* = 0.016; ipsilateral: *p* = 0.047, one-sided Wilcoxon signed-rank test; see Table S1 for detailed statistics). In contrast, while contralateral decoding in the BP task remained robust (*p* = 0.016), ipsilateral conditions did not reach significance (*p* = 0.078). Meanwhile, there was no difference in the noise ceiling levels between the two tasks (Figure S3). Binary decoding analyses for each object pair and the decoding time course also yielded similar results (Figure S5, S6).

To assess how spatial representations in the UP and BP tasks relate to perceptual encoding, we performed RSA, comparing each condition’s activation patterns to an independent sensory template (Figure S2). In the LOC (Figure 1c, right), contralateral representations showed significant correlations with the template in both UP (mean Fisher Z = 0.360, *p* = 0.016, Table S2) and BP (mean Fisher Z = 0.367, *p* = 0.016) tasks. By contrast, ipsilateral representations were only significant in the UP (mean Fisher Z = 0.328, *p* = 0.016), but not in the BP task (mean Fisher Z = 0.035, *p* = 0.281), indicating an absence of sensory-based ipsilateral coding under bilateral visual input.

In the EVC (Figure 1d), both decoding and RSA consistently revealed robust contralateral representation in the UP (decoding: *p* = 0.016; RSA: mean Fisher Z = 0.618, *p* = 0.016) and BP tasks (decoding: *p* = 0.016; RSA: mean Fisher Z = 0.676, *p* = 0.016). By contrast, ipsilateral decoding and RSA results failed to reach significance in either the UP (decoding: *p* = 0.891; RSA: mean Fisher Z = 0.039, *p* = 0.078) or the BP task (decoding: *p* = 0.5; RSA: mean Fisher Z = 0.082, *p* = 0.156). These findings suggest that ipsilateral representations in EVC are minimal and remain unaffected by changes in input configuration. This pattern is consistent with the strict retinotopic organization of early visual cortex (Tong et al., 2012; Wandell et al., 2007).

Collectively, these results demonstrate that ipsilateral representations in the LOC are robust under unilateral presentation but largely absent under bilateral conditions, consistent with previous findings (Bao & Tsao, 2018; Reithler et al., 2017). This reflects an adaptive spatial organization modulated by the visual input configuration, providing a baseline for the subsequent comparison with the VWM representation.

### VWM recruits bilateral LOC regions for both unilateral and bilateral object memory

Given the absence of ipsilateral LOC representations during bilateral perception (Figure 1e, left), we next asked whether similar spatial constraints govern VWM (Figure 1e, right). If VWM maintains the same perceptual representations, memorized object identity should remain lateralized. Alternatively, VWM might recruit additional ipsilateral areas to support a more distributed and bilateral memory code.

To test these two competing hypotheses, we designed two VWM tasks that differed in memory load. In the unilateral VWM task (retro-cue), two objects were presented bilaterally, followed by a spatial cue indicating which side’s object participants should maintain in memory (Figure 2a). Participants performed this task with high accuracy (mean accuracy = 0.955, *s.e.m.* = 0.012; Figure S7a), indicating strong task engagement.

**Figure 2.**
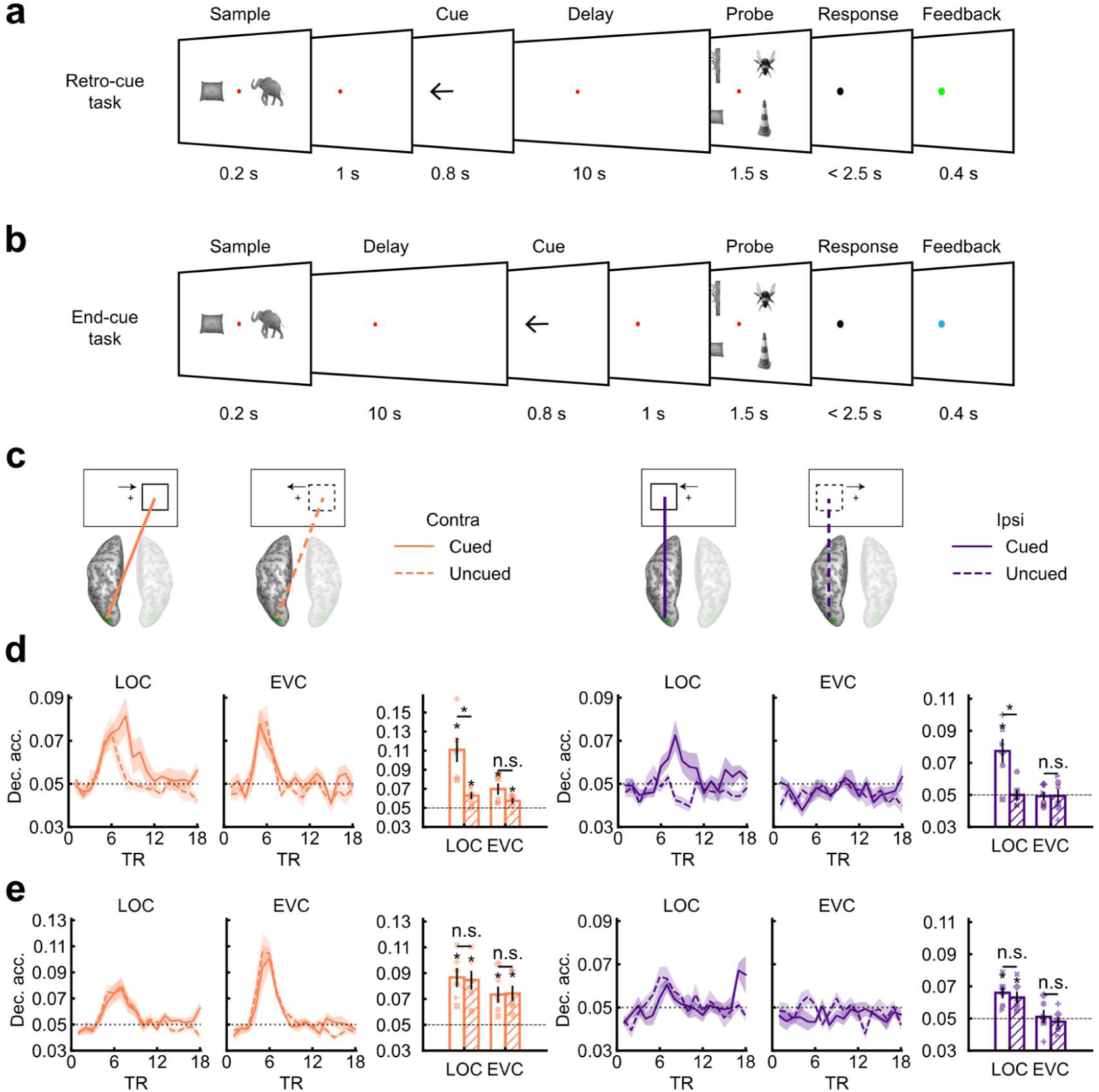
VWM task design and results. **a.** Illustration of the spatial retro-cue VWM task. Participants maintained fixation throughout the trial and memorized the cued object during the delay period. Afterward, they reported both the presence and identity of the cued object. **b.** Illustration of the spatial end-cue VWM task. Similar to **a.**, but participants had to retain both objects during the delay, as the cue was presented after the delay period. **c.** Schematic of decoding analysis using single hemisphere ROIs. Take the contralateral cued condition as an example (left diagram), for trials where the cued object was presented in the right visual field, only the BOLD signals from the left hemisphere ROIs were used for the decoding analysis (see Methods for details). **d.** Decoding accuracy under contralateral (left) and ipsilateral (right) conditions for the retro-cue VWM task. Line charts show the time course of decoding accuracy calculated using BOLD signals at each TR (TR = 1 s), while bar charts represent decoding accuracies computed using averaged BOLD signals across 6–10 TRs. In the bar charts, empty and dashed bars indicate cued and uncued conditions, respectively. **e.** Decoding accuracy under contralateral (left) and ipsilateral (right) conditions for the end-cue VWM task. The horizontal dashed black lines indicate chance level. Coral color represents the contralateral, while indigo color represents the ipsilateral condition. Shaded areas and error bars represent ± 1 *s.e.m.* Contra, contralateral; Ipsi, ipsilateral; Dec. acc., decoding accuracy; *s.e.m.*, standard error of the mean. * *p* < 0.05, uncorrected; n.s., non-significant, Wilcoxon signed-rank test.

We performed decoding analyses using BOLD signals at each TR, separating cued and uncued objects under contralateral and ipsilateral conditions (Figure 2c). As an example of this procedure, in the contralateral-cued condition, we decoded object identity using the ROI vertices in the opposite hemisphere. The same procedure was applied for ipsilateral-cued trials using the corresponding hemisphere (see Methods). Results from left- and right-field trials were averaged to generate the overall decoding time course (Figure 2d). For statistical comparisons, we also computed decoding accuracy based on the mean BOLD activity during the delay period (TRs 6–10; see Figure S7b for deconvolved responses).

As shown in the left bar chart of Figure 2d, contralateral decoding for cued objects in the LOC was significantly above chance and higher than for uncued objects (*p* = 0.016 for both comparisons, see Table S3, S4), confirming the critical involvement of LOC in object VWM. In contrast, although decoding accuracy for cued objects in EVC exceeded chance (*p* = 0.016), it did not differ from the uncued objects (*p* = 0.078), suggesting that EVC does not contribute to object VWM.

When decoding was performed using ipsilateral brain regions (Figure 2d, right), significant decoding was observed exclusively for cued objects in the LOC (cued vs. chance: *p* = 0.031; cued vs. uncued: *p* = 0.047), whereas decoding accuracy for uncued objects remained at chance (*p* = 0.719). In the EVC, neither cued nor uncued objects were decodable above chance (cued vs. chance: *p* = 0.719; uncued vs. chance: *p* = 0.656), and there was no difference in decoding accuracy between cued and uncued objects (*p* = 0.578). Similar results were obtained with binary decoding (Figure S8a–b, left).

These findings reveal that, unlike the contralateral-only representation observed in the BP task, object VWM recruits both contralateral and ipsilateral LOC. In contrast, neither hemisphere of EVC supports VWM representations. This suggests that the LOC plays an essential role in object VWM, with memory representations expanding to both hemispheres rather than being strictly constrained by perceptual configurations.

Would ipsilateral memory representations still emerge when participants must remember two objects presented in both visual fields? To address this question, we used a bilateral end-cue VWM task in which the spatial cue was presented only at the end of the trial (Figure 2b). In this modified setting, participants needed to retain both objects and their respective locations throughout the delay. Despite the increased memory load, participants still achieved high accuracy (mean accuracy = 0.947, *s.e.m.* = 0.008; Figure S7a). If ipsilateral decoding in LOC failed in this condition, it would suggest that such representations emerge only when a single object is prioritized, indicating task-dependent lateralization. Alternatively, successful decoding would support the notion that bilateral representations are a general property of object VWM.

In the end-cue task, object identity remained decodable in the contralateral LOC for both cued and uncued objects (Figure 2e left; cued vs. chance: *p* = 0.016, uncued vs. chance: *p* = 0.016, see Table S5, S6). More importantly, both cued and uncued objects were also successfully decoded under ipsilateral conditions (Figure 2e right; cued vs. chance level: *p* = 0.016; uncued vs. chance level: *p* = 0.016). No significant difference in decoding accuracy was observed between the two memorized items in either contralateral or ipsilateral condition (all *p*s > 0.28), indicating that the participants successfully memorized both objects simultaneously as the task required. Consistently, ipsilateral EVC showed no significant decoding for either cued or uncued objects (all *p*s > 0.5).

These results demonstrate that, unlike perception, VWM representations in the LOC are not strictly constrained by spatial lateralization. Instead, object information extends to ipsilateral regions, reflecting a more flexible coding scheme. Additionally, this ipsilateral recruitment persists even under high memory demands, indicating bilateral representation is a general feature of object VWM in the LOC.

### VWM extends ipsilateral LOC recruitment beyond perceptual constraints

Our previous analysis revealed that, although ipsilateral representation is minimal under bilateral input, VWM engages a robust ipsilateral representation, even under increased memory demands. However, the characteristics underlying this expansion remain unclear: (1) Do the additional ipsilateral regions recruited during VWM correspond to those that encode ipsilateral stimuli during perception, as in the UP task? (2) Does the extent of ipsilateral recruitment vary as a function of memory demand?

To address these questions, we conducted a vertex-ablation decoding analysis. First, we quantified each LOC vertex’s bilateral responsivity by measuring the similarity between its mean responses to 20 objects under contralateral and ipsilateral conditions in the UP task. Vertices with high bilateral responsivity were assumed to have a bilateral receptive field and to be inherently responsive to ipsilateral input during perception. We then ranked all vertices by this metric and progressively removed those with the highest values. After each removal step, we repeated the decoding analysis on the remaining (more unilateral) vertices. The resulting drop in decoding accuracy indicates the fraction of LOC vertices that support ipsilateral VWM representations (Figure 3a).

**Figure 3.**
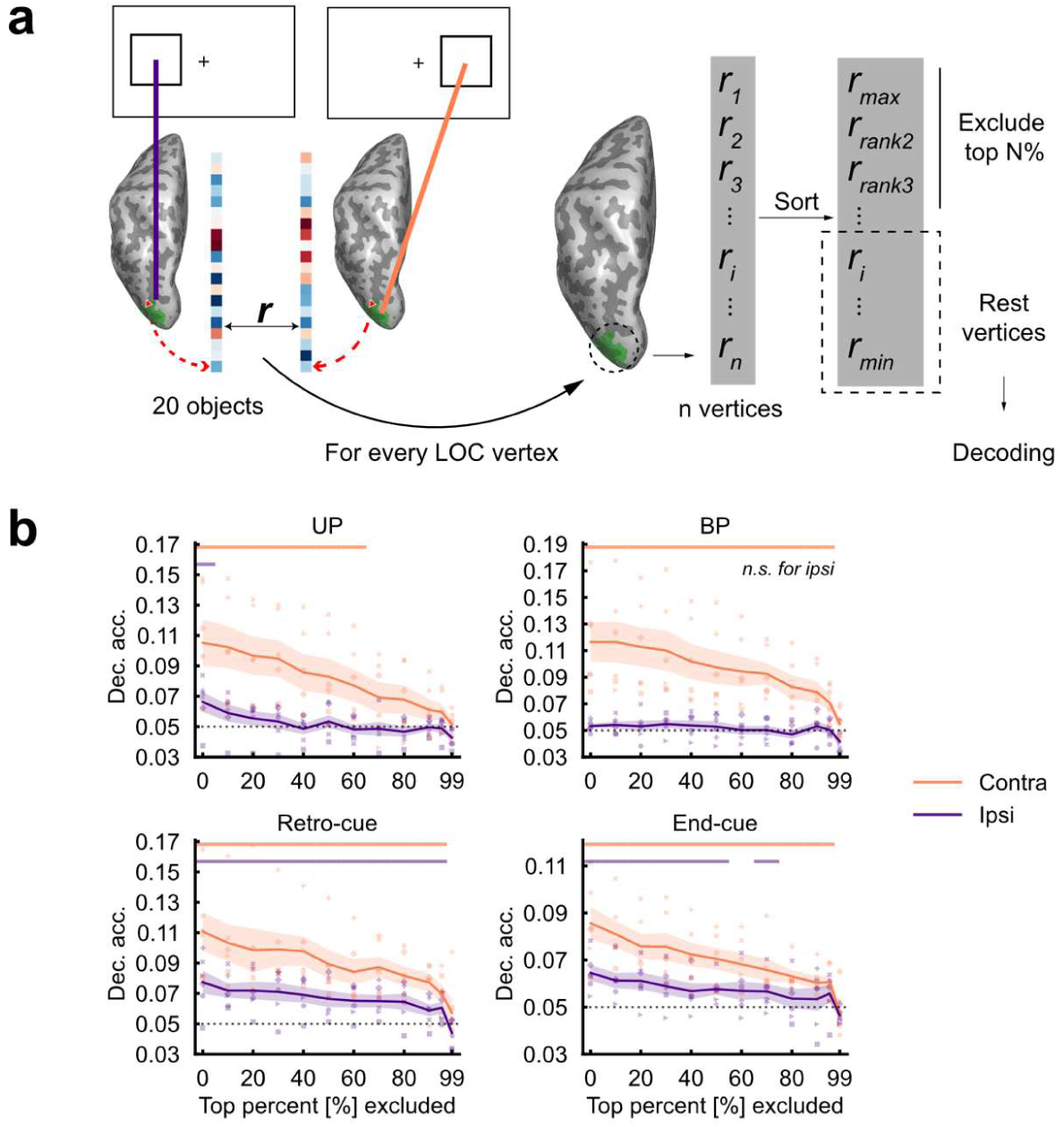
Vertex-ablation analysis in perception and VWM tasks. **a.** Schematic of the vertex-ablation analysis. For each LOC vertex, we calculated bilateral responsivity as the correlation between beta values from ipsilateral and contralateral conditions in the UP task. Vertices with the highest bilateral responsivity were progressively removed. **b.** Decoding accuracy in LOC following stepwise vertex-ablation analysis for the perception (first row) and VWM (second row) tasks. Each point along the curve represents decoding accuracy after excluding the top N% of vertices with the highest bilateral responsivity. For the VWM tasks, vertex-ablation analysis was conducted only on the cued condition. Horizontal colored lines above each figure indicated ablation steps where decoding was significant above chance (*p* < 0.05, uncorrected, Wilcoxon signed-rank test). The horizontal dashed black lines indicate chance level. Coral and indigo represent the contralateral and ipsilateral conditions, respectively. Shaded areas represent ± 1 *s.e.m*. Contra, contralateral; Ipsi, ipsilateral; Dec. acc., decoding accuracy; *s.e.m.*, standard error of the mean.

The ablation decoding results (Figure 3b) demonstrated that stable decoding of ipsilateral object information in the retro-cue VWM task persisted even after removing over 90% of the cortical vertices, with 40–80% of the vertices showing above-zero bilateral responsivity across participants (Figure S4). In the end-cue task, significant decoding remained after removing 70% of the LOC vertices, albeit to a less extent than in the retro-cue task (90%). In contrast, in either perception task, no significant ipsilateral representation was found after removing at most 10% of vertices.

These findings indicate that ipsilateral representations of objects in VWM expand to regions beyond what is observed in perception. VWM involves not only vertices inherently responsive to bilateral input, but also much of the ipsilateral LOC. Furthermore, increasing memory demands reduce the proportion of ipsilateral LOC regions involved for each object, highlighting the flexible recruitment of cortical areas according to task requirements.

### Increased attentional span plays a limited role in ipsilateral expansion

A potential confound is that the difference in attentional span between VWM and perception tasks may have contributed to our results, as an extended attentional range could also alter receptive field properties in the LOC (Kay et al., 2015). Could the extended ipsilateral representation observed in VWM simply reflect a wider attentional range during encoding, compared to the passive viewing condition in the perception task?

To address this concern, we conducted a bilateral attention control task (BA task, Figure 4a), where one of the objects rotated by a variable angle at random intervals. Participants pressed a key upon detecting a rotation in either visual field while maintaining central fixation. To ensure high attentional engagement, 90% of trials involved a rotation, and the rotation angle was adjusted via the QUEST procedure to ensure that overall accuracy converged to 70% (Watson & Pelli, 1983).

**Figure 4.**
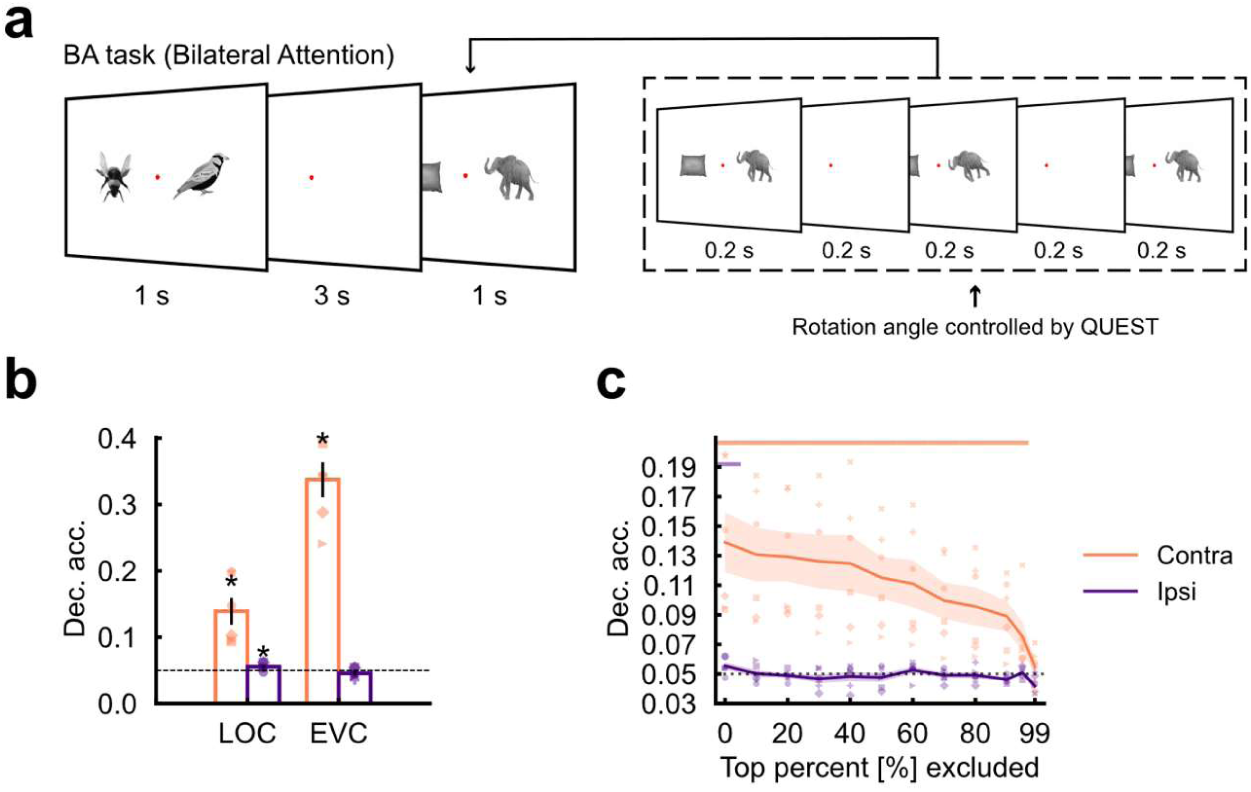
Attention task design and results. **a.** Illustration of the bilateral attention (BA) task. Similar to the BP task, two stimuli were presented three times in quick succession in each trial. Participants were required to detect the rotation of objects via a key press. **b.** Decoding accuracy for the contralateral and ipsilateral conditions in the LOC and EVC. **c.** Vertex-ablation decoding results for the BA task, analogous to Figure 3. Horizontal colored lines above indicated ablations steps where decoding was significant above chance (*p* < 0.05, uncorrected). The horizontal dashed black lines indicate chance level. Coral and indigo represent the contralateral and ipsilateral conditions, respectively. Shaded areas represent ± 1 *s.e.m*. Contra, contralateral; Ipsi, ipsilateral; Dec. acc., decoding accuracy; *s.e.m.*, standard error of the mean. * *p* < 0.05, uncorrected, Wilcoxon signed-rank test.

The results showed above-chance decoding accuracy in the ipsilateral LOC (Figure 4b, *p* = 0.031, see Table S7), with similar findings from binary decoding and decoding time course (Figure S5c–d, right; Figure S6). However, vertex-ablation decoding indicated that excluding 20% of LOC vertices resulted in chance-level decoding (Figure 4c). These results indicate that while an extended attentional span during encoding may contribute to ipsilateral recruitment, it cannot fully account for the broader ipsilateral representation observed in VWM.

### Cortical distribution of sensory-based ipsilateral representation

In the vertex-ablation analysis, we found that over 90% of LOC vertices contributed to object identity representation during the retro-cue VWM task, and over 70% during the end-cue VWM task—significantly more than those involved in perception and attention tasks. To visualize the cortical distribution of these representations across tasks, we employed an RSA-based searchlight analysis (Figure 5). Each participant’s LOC RDM from an independent central perception task (CP task, Figure S2) served as a common reference. For each task, we constructed an RDM for each vertex based on its neighboring vertices, and correlated these RDMs with the participant’s reference RDM. The resulting RSA maps were projected onto the common surface (fsaverage; Figure 5, S10). This analysis was applied to two VWM tasks (retro-cue task and end-cue task), as well as three additional tasks: unilateral object perception (UP), bilateral object perception (BP), and bilateral object attention (BA).

**Figure 5.**
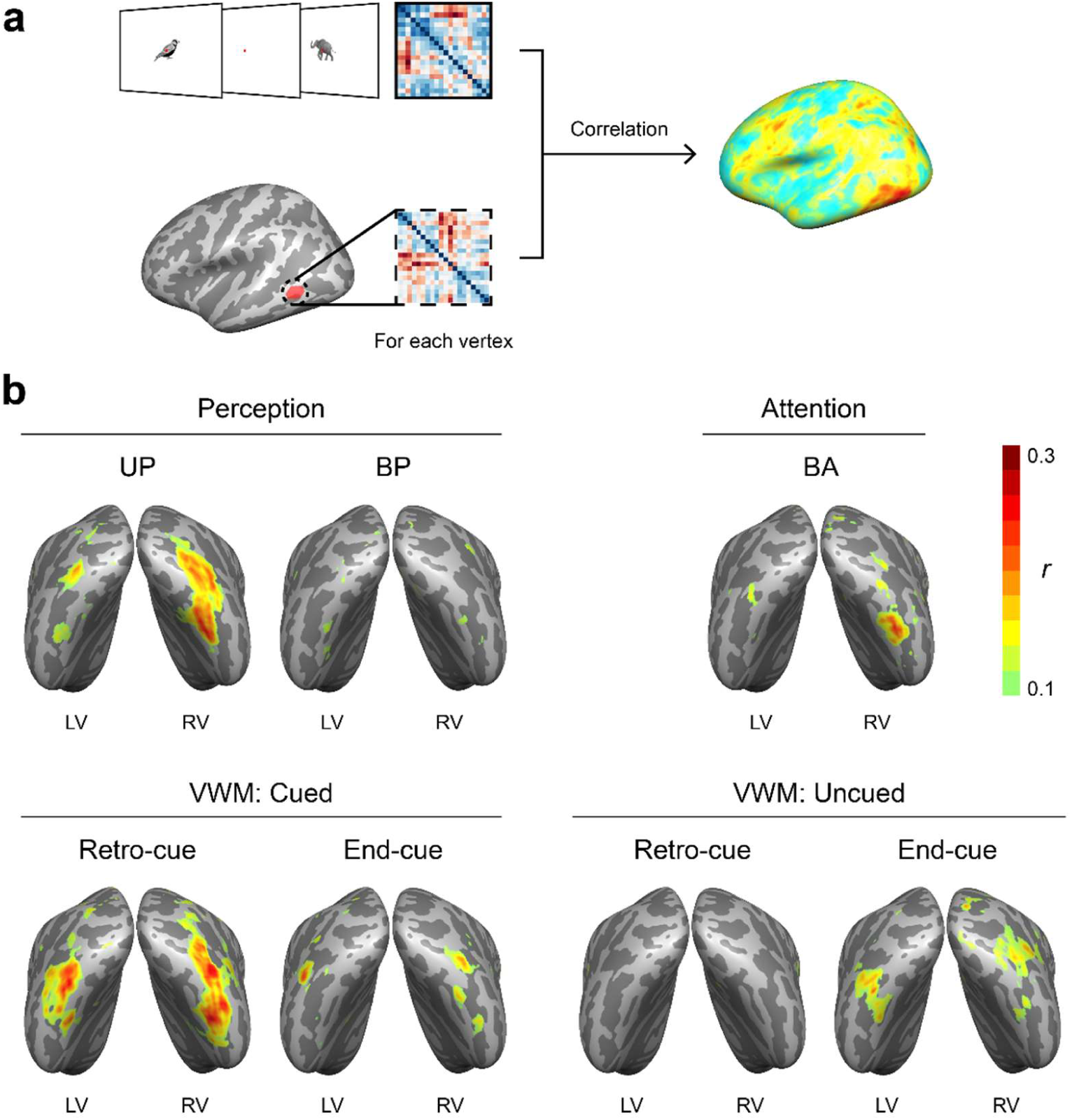
Cortical distribution of sensory-based ipsilateral representations. **a.** Schematic depiction of searchlight RSA. The RDM from the CP task in the LOC served as the reference RDM (Figure S2) and was correlated with floating RDMs from spatially localized searchlights in different tasks (see Methods). **b.** Ipsilateral representations revealed by searchlight RSA across perception, attention and VWM tasks. Only vertices with Pearson *r* values > 0.1 are shown. “LV” (left visual field) and “RV” (right visual field) labels indicate that values on each hemisphere were calculated based on the stimulus label in the corresponding visual field. For the left hemisphere, values correspond to the LH–LV sub-condition (left hemisphere activation for left visual field stimuli); for the right hemisphere, values correspond to the RH–RV sub-condition (right hemisphere activation for right visual field stimuli).

Focusing on cortical distribution of the representation to the object presented in the same-side visual field, we observed strong ipsilateral representations during the UP task, but only limited ipsilateral engagement in the BP and BA tasks (Figure 5b, top). This was supported by higher averaged correlation values among LOC vertices in the UP compared to BP and BA (Figure S11; UP vs. BP: *p* = 0.016; UP vs. BA: *p* = 0.016; see Table S8).

In VWM, the retro-cue task exhibited a broader ipsilateral representation for cued objects relative to uncued objects (Figure 5b, bottom left), showing higher mean correlation values (Figure S11; *p* = 0.016). This enhanced similarity for cued objects in retro-cue task was also greater than in the BP and BA tasks (retro-cue cued vs. BP: *p* = 0.016; retro-cue cued vs. BA: *p* = 0.016), but not significantly different from the UP task (*p* = 0.281). The end-cue task also revealed sensory-based ipsilateral representation (Figure 5b, bottom right), as evidenced by higher correlation than the BP task (cued: *p* = 0.031; uncued: *p* = 0.031). However, this representation was more expanded in the retro-cue task than in the end-cue task (retro-cue cued vs. end-cue cued: *p* = 0.016; retro-cue cued vs. end-cue uncued: *p* = 0.016), possibly due to the need to divide resources between both objects in the end-cue task (end-cue cued vs. end-cue uncued: *p* = 0.578).

Together, our results show that VWM is associated with both the expansion and spatial redistribution of sensory-like ipsilateral coding across the LOC. This finding indicates a fundamental reorganization of representational topology alongside memory flexibility, beyond what is captured by global ablation metrics alone.

### Divergent spatial representation strategies in association areas during VWM

In addition to high-level visual regions, association areas, such as IPS and PFC, may also play a critical role during VWM (Yu & Shim, 2017), with recent studies suggesting that the spatial constrains of VWM extend to these areas as well (Brincat et al., 2021). To test whether association regions show similar spatial expansion of memory representations as the LOC, we conducted decoding analysis in the IPS and PFC in our VWM tasks.

First, our results revealed robust memory-specific effects in both IPS and PFC during the retro-cued task: significant decoding was observed for cued versus uncued objects in both contralateral (IPS: *p* = 0.016; PFC: *p* = 0.031, see Figure 6a left and Table S4) and ipsilateral (IPS: *p* = 0.016; PFC: *p* = 0.031, Figure 6b left) conditions. These results show that association areas contribute to maintaining mnemonic representations and that, like the LOC, both IPS and PFC show expanded spatial representations during the retro-cue VWM.

**Figure 6.**
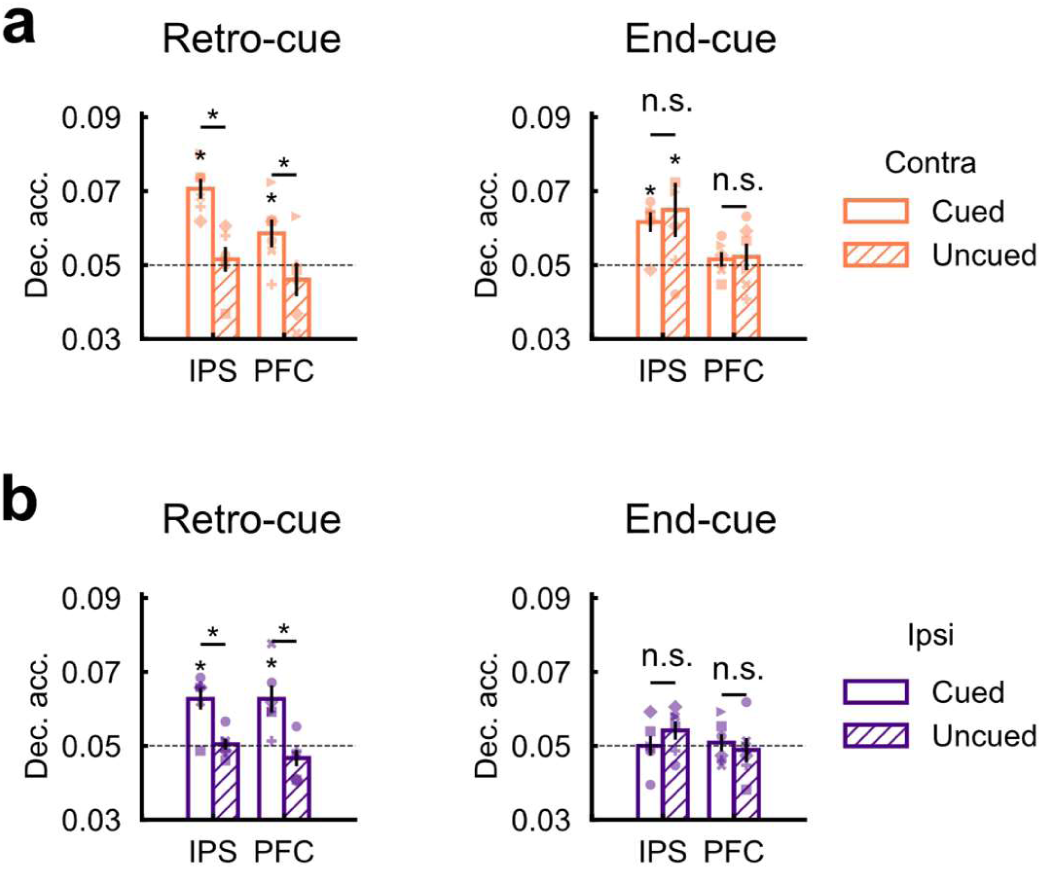
Decoding accuracy in association areas in VWM tasks. Decoding accuracy in the IPS and PFC during retro-cue (left) and end-cue (right) VWM tasks. **a.** Contralateral decoding results. **b.** Ipsilateral decoding results. The horizontal dashed black lines indicate chance level. Error bars represent ± 1 *s.e.m*. Contra, contralateral; Ipsi, ipsilateral; Dec. acc., decoding accuracy; *s.e.m.*, standard error of the mean. * *p* < 0.05, uncorrected, n.s., non-significant, Wilcoxon signed-rank test.

However, in the end-cue task, while the contralateral IPS continued to represent object identity for both the cued (*p* = 0.031, see Figure 6a right and Table S5) and uncued (*p* = 0.047) conditions, no ipsilateral representation was detected (Cued: *p* = 0.656; Uncued: *p* = 0.109, Figure 6b right). Moreover, the PFC showed no significant object-identity representations under any condition in this task (all *p*s > 0.34).

These results indicate that the IPS adopts a global, expanded object representation strategy when a single object is maintained, but reverts to a location-constrained code when multiple items are held. In contrast, the PFC shows a representation expansion only for single-object memory, and under high load, does not exhibit either expansion or location specificity. Instead, it might adopt a more abstract, non-linear coding format, as reflected by the negative decoding results for separate object identity when two objects were maintained.

We next asked whether association areas could be the main source of the ipsilateral representation in the LOC via top-down feedback. To test this, we performed cross-region RSA for both VWM tasks, comparing the representational patterns of ipsilateral LOC with contralateral LOC and other regions. Using partial regression to isolate the unique contributions of each region’s RDM in each condition, we found that ipsilateral LOC representations correlated most strongly with contralateral LOC, more so than with IPS or PFC (Figure S9). This indicates that ipsilateral representations in the LOC likely mirror contralateral patterns and are not primarily driven by feedback from association areas.

Collectively, these results echo previous findings highlighting the involvement of association areas in maintaining mnemonic information (Xu, 2023; Yu & Shim, 2017; Bettencourt & Xu, 2016). However, top-down feedback from the IPS and PFC might not be the primary driver of the expanded ipsilateral representation in high-level visual areas. Instead, association and visual regions diverge in their spatial coding strategies, with this divergence becoming more pronounced under high memory load.

## Discussion

The present study reveals a fundamental divergence in the spatial organization of object representations between visual perception and VWM within the LOC. While perceptual representations in the LOC are spatially constrained by visual field occupancy, VWM recruits ipsilateral cortical regions beyond those engaged during perception and attention. Moreover, this expanded representation persists across varying memory demands. Although the IPS and PFC contribute to memory maintenance, our findings suggest that top-down feedback from these association areas is not the primary driver of the expanded ipsilateral representation in high-level visual areas. Together, these results highlight the existence of spatial reorganization in high-visual cortex during VWM compared to perception, revealing a more flexible and globally distributed coding strategy for VWM.

Rather than being highly invariant and stable, LOC representations are susceptible to external stimulus configurations and internal cognitive demands during perception. Previous fMRI studies have reported bilateral representations in LOC under unilateral perception paradigms (Silson et al., 2022; Reithler et al., 2017; Cichy et al., 2013, 2011), likely due to its large receptive fields encompassing both visual hemifields (Kravitz et al., 2008). Our findings support this view and further demonstrate that during bilateral object presentation, object coding in LOC is largely confined to the contralateral hemisphere, likely reflecting competitive interactions that prioritize contralateral input. However, this spatial constraint is overcome during VWM, where robust bilateral representations persist even when maintaining two objects simultaneously. This suggests that VWM employs a more flexible spatial organization, adapting to memory demands rather than simply replicating the perceptual coding.

A key finding of this study is that VWM-related spatial expansion is selective to LOC rather than universally distributed across the visual hierarchy. While EVC can exhibit ipsilateral representations when retaining low-level features (Zhao et al., 2021; Pratte & Tong, 2014; Ester et al., 2009), our results indicate that high-level object stimuli solely engage ipsilateral LOC during VWM, despite EVC also contributing to object perception (Figure 1d). This raises an important question: Is memory-related spatial expansion driven by stimulus complexity or task relevance (Teng & Postle, 2021; Pratte & Tong, 2014; Ester et al., 2009)? Future studies should systematically manipulate different levels of memory requirements to determine whether VWM for real-world objects engages EVC in a similar bilateral manner.

Understanding the mechanisms underlying the expanded spatial representation in VWM remains a critical question. One possibility is that this expansion arises from changes in population receptive field (pRF) properties during encoding. VWM requires a broader attentional span than perception, which could extend receptive fields and allow ipsilateral encoding (Kay et al., 2015). However, our data suggest that such attentional modulation alone is insufficient to account for the broad ipsilateral representations observed in VWM (Figure 4), implying additional underlying mechanisms. Another possibility is that ipsilateral representations arise via top-down feedback from association cortices such as the parietal and prefrontal regions (Zhao et al., 2021). However, our cross-region RSA also argues against a purely feedback-driven mechanism (Figure 6), as ipsilateral LOC activity closely mirrored contralateral sensory patterns rather than abstracted representations from IPS or PFC. Instead, intrinsic visual network dynamics, particularly interhemispheric coordination and communication between homologous LOC regions, may drive this spatial expansion (Robinson et al., 2025; Zhao et al., 2021). While our RSA analysis aligns with this hypothesis (Figure S9), the limited temporal resolution of fMRI prevents direct causal inferences. Future studies could address this by leveraging high-temporal-resolution methods, such as high-density electrophysiological recordings or intracranial EEG, to simultaneously record bilateral IT cortex activity during memory tasks in primates, thereby clarifying the mechanisms of interhemispheric information communication.

Another important open question is when such bilateral expansion during VWM occurs. While fMRI cannot resolve these temporal dynamics, EEG studies provide relevant insights. Previous studies have shown a strong contralateral bias during encoding, but equally strong decoding from bilateral posterior channels during VWM reactivation (Wolff et al., 2020), suggesting a post-encoding shift toward bilateral memory traces over time. A relevant phenomenon that may be related to the emerging reorganization comes from the contralateral delay activity (CDA) in EEG. The difference between contralateral and ipsilateral responses relative to the visual field of the remembered items reflects VWM storage (Wang et al., 2019). Interestingly, the contralateral-ipsilateral differences gradually reduced during the delay period, as observed in both CDA amplitude reductions (Vogel & Machizawa, 2004), and declines in the decoding accuracy based on the bilateral response difference (Fukuda et al., 2016). These findings suggest that VWM reorganizes object representations spatially, shifting from contralateral dominance to bilateral encoding during the delay period. However, it remains unclear whether this shift reflects a weakening of the strong contralateral sensory input or the construction of a new memory trace, as well as where in the cortex this reorganization takes place, given the spatial limitations of EEG. Further computational modeling and precise neural recordings are needed to resolve this question.

Although we observed spatially expanded representations in VWM, other studies have reported task-dependent spatial constraints on memory representations. In early visual areas, bilateral representations are observed when a single item is remembered without a location requirement (Ester et al., 2009), but when both visual fields are task-relevant, representations become more spatially constrained (Pratte & Tong, 2014). Behavioral studies support this task-dependent flexibility, showing that when participants encode orientations across both visual fields, the stored feature can modulate perception in a spatially specific manner. However, when only one grating is remembered, its influence extends more globally, unless spatial specificity is reinforced (Teng & Postle, 2021). Our end-cue and retro-cue VWM tasks consistently revealed bilateral representations, regardless of spatial memory demands. This suggests that object VWM inherently favors global representations in the LOC, independent of spatial precision requirements—a pattern that may differ from early visual cortices.

In summary, our findings reveal a spatially extended, sensory-based VWM representation in object-selective regions of the LOC. Unlike perception, which is constrained by spatial organization, VWM flexibly expands its representational range based on input configuration, task modality, and memory demands. Future studies should employ EEG and MEG to pinpoint the precise temporal dynamics of bilateral representation emergence in VWM. Additionally, high-throughput electrophysiological recordings in non-human primates could provide high-temporal-resolution insights into interhemispheric coordination in object-selective cortex at the single-unit level. Comparing different probing methods may also clarify their influence on VWM representational structure. Together, these approaches will advance our understanding of the neural mechanisms driving the spatial expansion of VWM and its interactions with perception, attention, and executive function.

## Methods

### Participants

Six healthy participants (three males, age range 21-29 years, mean age = 23.0 years, *s.e.m.* = 1.13) underwent all experiments in the fMRI scanner. All participants had normal or corrected-to-normal vision, were right-handed, and reported no history of psychiatric or neurological disorders or family history of such conditions. None had used psychotropic medications in the past six months. The sample size followed previous VWM decoding studies that have demonstrated reliable effects with cohorts of comparable size (Zhao et al., 2021; Harrison & Tong, 2009). Crucially, an extensive amount of fMRI data was collected from each participant, consisting of six different tasks across more than 12 scanning sessions, containing 1520 event-related VWM trials. This sampling design maximizes statistical power for within-subject comparisons and allows us to probe general object representation properties while maintaining identical stimulus sets across participants. The resulting rich data per participant provide a rigorous foundation for task-level contrasts and support the generalizability of our conclusions despite the modest cohort size. This study was approved by the Ethics Committee of School of Psychological and Cognitive Sciences at Peking University (Project ID: 2022-03-04). Written informed consent was obtained from each participant prior to the experiment, and they were compensated for their participation.

### Stimuli and Experimental Procedure

All six participants completed six experimental tasks, including three perception tasks, two VWM tasks, and one attention task over 12–16 separate scanning sessions. Each scanning session lasted 0.75–1.5 hours. Additionally, two localizer tasks were performed prior to the experimental tasks. All experiments were generated via PsychToolBox-3 (Kleiner et al., 2007; Brainard, 1997; Pelli, 1997) on MATLAB 2016b (MathWorks).

Across all tasks, 20 grayscale images of isolated real-world objects were used as stimuli (Figure S1a). In the perception and attention tasks, stimuli were presented at a visual angle of 10°, employing a fast-event design. In the VWM tasks, stimuli appeared at 10° visual angle in the encoding array and 5° visual angle in the probe array, employing an event-related design. All stimuli were presented on a white background. Two display systems were used across participants and tasks: one back-projected onto a translucent screen inside the scanner, and the other displayed on an LCD screen outside the scanner bore. Both systems operated at a refresh rate of 60 Hz and a spatial resolution of 1024 x 768 pixels. Stimulus presentation programs were adjusted to ensure consistent visual angle across both display systems. During scanning, participants lay supine and viewed the stimuli via a mirror mounted on the head coil.

#### UP task

Unilateral Perception task (Figure 1a). In each trial, a single object was presented on one side of the screen, with a 0.3° offset between the edge of the image and the center. The object was flashed three times within 1 second (200 ms on, 200 ms off), followed by a 3-second blank. Each object was presented 20 times per side, resulting in 800 valid trials across 10 scanning runs. Each run consisted of 100 trials, including 80 valid trials and 20 empty trials, each lasting 4 seconds. The lead-in and lead-out times were both set to 6 seconds, resulting in a total run duration of 412 seconds per run. Within each scanning run, participants were required to perform a fixation task, with the central fixation point alternating between “+” and “×” every 3– 7 seconds. Participants were instructed to respond with a key press upon detecting a fixation change.

#### BP task

Bilateral Perception task (Figure 1b). The presentation settings and task requirements were similar to the UP task, except that two objects were presented simultaneously. Each object appeared on each side of the screen, with a 0.3° gap between each object’s edge and the central fixation. All possible pairings of the 20 objects yielded 400 valid trials across 4 scanning runs. Data from 20 valid trials where the same object appeared in both visual fields were discarded from analysis. Each run contained 100 valid trials and 25 empty trials, each lasting 4 seconds, resulting in a total run duration of 512 seconds. Trials in which the same object appeared in both visual fields (*n* = 20) were excluded from analysis.

#### CP task

Central perception task (Figure S2a). The presentation settings were similar to the UP task, except that the objects were presented centrally. In each trial, the object was flashed three times, and in 10% of the trials, the contrast of the non-white pixels in the object was reduced by 50% during the middle flash. Instead of using a central fixation task, participants were instructed to press a key upon detecting the contrast change in the object. Each object was presented 15 times, yielding 300 valid trials across 3 runs. Each run contained 100 valid trials and 25 empty trials, each lasting 4 seconds, resulting in a total run duration of 512 seconds.

#### VWM retro-cue task

As shown in Figure 2a, the VWM task was a modified retro-cue working memory task based on previous studies (Pratte & Tong, 2014; Harrison & Tong, 2009), incorporating a spatial retro-cue rather than a sequential one. In each trial, participants viewed the encoding array consisting of two objects simultaneously presented in the left and right visual fields for 0.2 seconds, with a 0.3° offset between the edge of the image and the center (same as in the BP task). This was followed by a 2.25° visual angle arrow cue presented at the center indicating the object to be memorized. After a 10-second delay, a probe array containing four objects was presented, with the images offset by 2.5° from the horizontal and vertical midlines. Each position of the probe objects corresponded to a button on the response box, and participants were instructed to select the corresponding object via button press or withhold response if the cued object was absent. Feedback regarding accuracy of the current response was provided by a fixation color change following the button press.

The objects in the encoding array were composed of two different items selected from the 20 objects. To counterbalance the number of presentations of different objects, spatial locations, and cueing positions, all possible permutations were generated, ensuring that each of the 20 objects appeared an equal number of times as the cued or uncued object in both visual fields. This yielded a total of 760 trials per participant, with 19 trials for each object per condition:

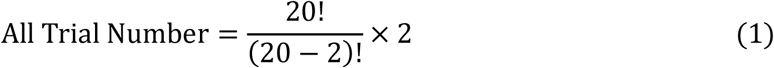

As for the probe array, in 75% of the trials, the cued object appeared at one of the four possible screen positions, each with approximately equal probability (142–144 trials per position) to prevent response biases. In the remaining 25% of the trials, the cued object was absent. The other objects in the probe array were randomly selected from a subset of 100 objects, which were manually selected from the same larger image set as the 20 objects (Bao et al., 2020). All four objects in the probe array were unique for a certain trial. Each run consisted of 20 trials, each lasting 16 seconds, with inter-trial-intervals (ITIs) of 2, 4, 6, 8, or 10 seconds randomly distributed across trials. A 6-second lead-in period was included at the beginning of each run, with no lead-out time, resulting in a total run duration of 446 seconds. Participants completed 38 scanning runs across 5–6 scanning sessions.

#### VWM end-cue task

As shown in Figure 2b, the VWM end-cue task was modified from the retro-cue task described above, differing only in the appearing time of the cue. We moved the cue to the end of the delay period, requiring participants to retain both objects throughout the delay. The trial settings were similar to the previous retro-cue VWM task, with all object orders and combinations newly generated. This task also comprised 38 runs, each lasting 446 seconds, distributed across 4–6 scanning sessions for each participant to finish.

#### BA task

Bilateral attention task (Figure 4a). The attention task was adapted from the BP task, with the following modifications to increase attentional demands. Instead of a changing fixation point, a constant red dot was presented at the center. In each trial, objects also flashed three times, and during the middle frame, one object on either side rotated slightly. The rotation angle was controlled by the QUEST procedure (Watson & Pelli, 1983) to converge on 70% accuracy. Participants needed to press one key for a leftward rotation and another key for a rightward rotation. Each run contained 100 valid trials and 25 empty trials, each lasting 4 seconds, resulting in a total run duration of 512 seconds. Data from 20 valid trials where the same object appeared in both visual fields were discarded from analysis.

### Localizer tasks

#### Retinotopic mapping

To map early visual cortex regions for each participant, we conducted the population receptive field (pRF) experiment and estimated pRF as described by previous studies (Benson et al., 2018; Kay et al., 2013). Throughout the experiment, participants were required to perform a fixation color judgment task, pressing a button whenever they detected a change in color. Stimuli consisted of wedge, ring, and bar apertures composed of objects scattered on a pink-noise background. In the first two runs, wedges rotated clockwise and counterclockwise around the fixation point; in the next two runs, rings expanded outward or contracted inward from the fixation point; and in the final two runs, bars moved along the diameter with different directions. Stimuli subtended 16° of visual angle. Each pRF run lasted 300 seconds, with the aperture and images updating at 15 Hz.

#### High-level visual areas localization

To define high-level visual regions, we used a category-selective localizer with images of bodies, faces, scenes, objects, words, and Fourier phase-scrambled images. The localizer task contained two block-design runs, each lasting 396 seconds and containing 24 blocks (6 categories × 4 repeats). The lead-in and lead-out times were both set to 6 seconds. Each block lasted 16 seconds. Within each block, 16 images from the same category were presented for 500 ms followed by a 500 ms blank interval. Isolated objects or scenes were overlaid on a scrambled background and displayed centrally at a visual angle of 8°. Participants performed a 1-back task, pressing a button when the current image matched the previous one to stay engaged with the task.

### fMRI data acquisition and preprocessing

All fMRI data were collected using a 3 T Siemens Prisma MRI scanner at the Center for MRI Research at Peking University, equipped with a 64-channel phased-array head coil. In each scanning session, a high-resolution T1-weighted anatomical image was acquired using a 3D-MPRAGE sequence for cross-session registration. Functional images during all tasks were acquired using a T2*-weighted echo-planar imaging (EPI) sequence, with the following parameters: TE = 30 ms, TR = 1000 ms, flip angle = 78°, voxel size = 2.5×2.5×2.5 mm, slice thickness = 2.5 mm, total slices = 56. Field maps were also acquired prior to all other scans in each session.

The data were preprocessed using BrainVoyager QX software (version 2.8; Brain Innovations, Maastricht, The Netherlands) and custom MATLAB code in conjunction with the NeuroElf toolbox (https://neuroelf.net/). The anatomical image from the first scanning session of each participant was manually aligned to the AC–PC line and transformed into the Talairach space. All subsequent T1 images were aligned to the first Talairach-transformed structural image. Cortical surfaces were reconstructed by segmenting gray and white matter, followed by inflation of the white/grey matter boundary.

Functional images were initially corrected for EPI distortions using the field maps via the anatabacus plugin in BrainVoyager QX. Subsequent preprocessing included temporal correction, motion correction (trilinear/sinc interpolation), spatial smoothing (FWHM = 4 mm), and high-pass temporal filtering (GLM-Fourier with 2 sine/cosine pairs). The functional images closest in time to the T1 scan were used as a reference and aligned to the T1 image, and all other functional images were aligned to this reference. The first six TRs of each run were discarded to minimize the transient magnetic saturation effect. Next, BOLD signals were extracted from all the remaining data for each scanning run. For each vertex, the signals were normalized across all remaining TRs.

In order to project the results on a common surface space in the searchlight RSA, we used fMRIprep 22.0.1 (Esteban et al., 2019) to re-preprocess all the data. Structural MRI images underwent intensity correction and skull stripping, and were then normalized to the MNI152NLin2009cAsym template. Brain tissue segmentation into CSF, WM, and GM was performed using FSL, and brain surfaces were reconstructed with FreeSurfer. For functional MRI, each BOLD run was motion corrected, slice-time corrected and then co-registered to the T1w reference. The data were then resampled to both standard and surface spaces using ANTs and FreeSurfer. Internal operations leveraged Nilearn for functional processing. All preprocessed data were transformed to BrainVoyager files via fp2bv toolbox (https://github.com/UW-CHN/fp2bv).

For the perception and attention tasks, a general linear model (GLM) was used to estimate the beta responses for each trial via custom MATLAB scripts. For the VWM tasks and retinotopic tasks, raw BOLD responses were used for analysis. Data in each run were normalized across TRs for each vertex. For the block-design localizer task, we used voxel-wise GLMs within BrainVoyager QX software to calculate the contrast maps, and these volumetric maps were subsequently projected onto the individual cortical surface.

### Regions of interest (ROI) definitions

All ROIs were defined on the participant-specific surface space by combining functional activation with an established anatomical atlas.

For high-level visual areas, the lateral occipital complex (LOC) was defined using data from the localizer task as described above. A contrast of [face + body + object + scene – 4 × scrambled] was used to define the LOC. The positive regions near the lateral occipital areas on the contrast map were selected, with a minimum statistical threshold of *q*(FDR) < 0.05 for each participant.

We used data from retinotopic mapping to localize early visual areas. The pRF estimation was calculated using scripts provided by Benson et al. (2018) and Kay et al. (2013). Based on the pRF results, we defined V1–V3 as in Benson et al. (2018). These regions were then combined together to form the early visual cortex (EVC) ROI.

For association areas, the intraparietal sulcus (IPS) and the prefrontal cortex (PFC) were defined by combining VWM-evoked activation with atlas-based anatomical constraints. Specifically, whole-brain VWM activations during the retro-cue and end-cue tasks were calculated by fitting voxel-wise GLMs to each run, using regressors for sample-array and probe-array onsets convolved with a canonical hemodynamic response function. The resulting beta contrast maps were thresholded at *q*(FDR) < 0.05 to obtain the VWM-evoked regions. The suprathreshold voxels from both tasks were combined to produce a single VWM activation mask for each participant.

The anatomical position of the IPS was defined using the visual topography probabilistic maps from Wang et al (2015). Probability maps for IPS0-IPS4, originally in MNI space, were transformed to each participant’s Talairach space and projected onto their cortical surface. Vertices belonging to these subregions with a probability > 20% were combined to form a single anatomical IPS mask. The final IPS ROI for each participant was defined as the overlap between this anatomical mask and the participant’s VWM activation map.

The anatomical position of the PFC was localized using the HCP-MMP1.0 atlas (Glasser et al., 2016; Horn, 2016). Atlas parcels corresponding to dorsolateral prefrontal regions (areas 8C, 8Av, i6-8, s6-8, SFL, 8BL, 9p, 9a, 8Ad, p9-46v, a9-46v, 46, and 9-46d) were transformed from MNI to Talairach space and mapped onto each participant’s surface. These parcels were merged into one anatomical PFC mask, which was then intersected with the participant’s VWM activation map to yield the final PFC ROI.

### Multivariate Pattern Analysis

We used decoding and representational similarity analysis (RSA) to investigate the neural representational patterns. In VWM tasks, BOLD time series data were extracted from each ROI and segmented into 18-second segments based on trial onset. In the perceptual and attention tasks, beta values from each trial were used. All trials within each condition were used as samples, with vertex-wise BOLD or beta values within each ROI serving as features for subsequent analysis.

#### Decoding analysis

In the main decoding analysis, a one-vs-rest multiclass classifier (20 classes) was constructed using a linear SVM algorithm based on Python’s scikit-learn library. Leave-one-trial-out cross-validation was used, training the model on n−1 samples and testing on the held-out sample. Classification accuracies from the n folds were averaged to obtain the final decoding accuracy (chance level = 0.05). In addition, for each task condition, pairwise binary decoding was performed for all object pairs. The resulting decoding accuracies were used to construct a decoding-based RDM. The averaged decoding accuracy across all objects was calculated using the averaged off-diagonal values in the RDM (chance level = 0.5).

In all displayed results, decoding accuracies were shown separately for the contralateral and ipsilateral conditions. Specifically, in perception and attention tasks, the decoding accuracy for the contralateral and ipsilateral conditions was calculated as:

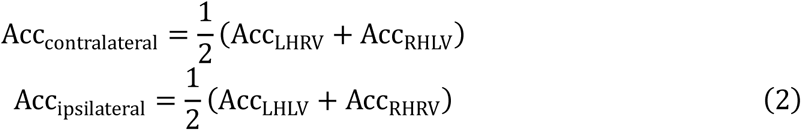

where LHRV refers to decoding object labels in the right visual field (RV) using beta values from the left hemisphere (LH) ROI, and RHLV refers to decoding object labels in the left visual field (LV) using beta values of each trial from the right hemisphere (RH) ROI. Similarly, LHLV and RHRV correspond to decoding object labels within the same-side hemisphere and visual field (i.e., ipsilateral conditions).

For VWM tasks, the full set of 760 trials was split into two sets, each with 380 trials based on cue direction (left or right). Then, the decoding accuracy for each condition was calculated as:

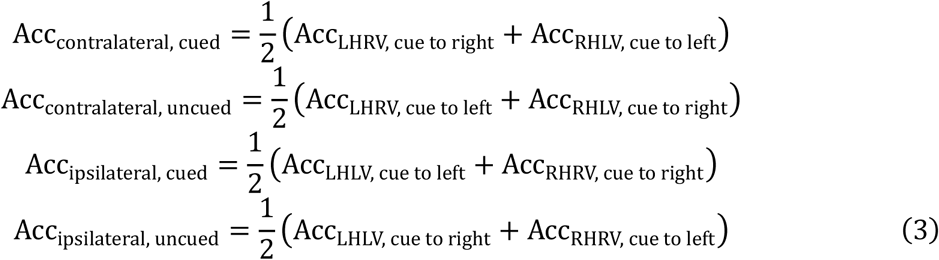

where the LHRV and RHLV conditions represent decoding object identity in the right and left visual fields using left and right hemisphere ROIs, respectively. Each sub-condition (e.g., LHRV, cue right) included 380 out of 760 trials with the cue directed to the relevant side. For example, in the *contralateral, cued* condition, the LHRV sub-condition used BOLD signals from the left hemisphere ROI to decode object identity in the right visual field during trials cued to the right. Similarly, the RHLV sub-condition used the other 380 trials cued to the left, with right hemisphere ROI BOLD signals decoding the object label in the left visual field. The decoding accuracy for the *contralateral, cued* condition was computed as the average of these two sub-conditions.

#### Vertex-ablation analysis

To assess the proportion of LOC vertices contributing to object identity representation, we performed a vertex-ablation analysis using beta values from the BP task. The goal was to isolate and exclude vertices with similar response profiles across the two visual fields, which may reflect inherently bilateral visual input. For each vertex in the LOC, the correlation between its beta values for the 20 objects in the ipsilateral and contralateral conditions was calculated and served as the bilateral responsivity:

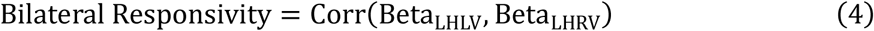

We performed stepwise vertex-ablation by progressively excluding the vertices with the highest bilateral responsivity for both the contralateral and ipsilateral condition. At each step, an increasing percentage of vertices (N%) were excluded, and decoding accuracy was recalculated based on the remaining vertices.

#### Representational similarity analysis (RSA)

To test for the representational pattern similarities across tasks, we conducted RSA based on the task-specific representational dissimilarity matrices (RDMs). For each condition, we first constructed the RDM by averaging the beta values across all trials for each object label, then computed the pairwise Pearson correlation *R*(object A, object B) between every object pair using the beta values from the ROI vertices. The dissimilarity between object pairs was computed as 1-*R*(object A, object B), resulting in a 20 × 20 symmetric RDM for each condition. For the CP task, RDMs were calculated by combining beta values from ROIs in both hemispheres; for all other tasks, RDMs were calculated using beta values from a single hemisphere ROI. RDMs in different conditions were calculated based on the data and labels described above. Representational similarity was quantified as the Pearson correlation between the off-diagonal values of two corresponding RDMs. Typically, each condition’s RDM was compared against the CP task RDM, which served as a common sensory template. Contralateral and ipsilateral RSA results were obtained by averaging these correlations across sub-conditions (formula 2–3). All results were calculated at the individual level and Fisher Z-transformed prior to group-level statistical comparisons.

#### Searchlight-based RSA

Searchlight RSA was performed in both participants’ native surface space and the fsaverage space. We first constructed reference RDMs and floating RDMs. The reference RDMs were derived from the LOC in the CP task (Figure S2), where objects were presented at the center. Each floating RDM was calculated for each vertex using the BOLD or beta values from the 200 surrounding neighbor vertices in the native surface space, and 400 surrounding neighbor vertices in the fsaverage space. Neighbor vertices numbers of 100, 200, and 300 were also tested and did not produce systematic differences from the displayed results. In the two VWM tasks, we used the average BOLD signal from the 6–10 TRs for constructing VWM RDMs; in all other tasks, trial-wise beta values were used. To assess representational similarity for each vertex, we computed the Pearson correlation between the off-diagonal values of the reference RDM and each floating RDM. In Figure 5, we present the representational similarity for ipsilateral condition, comparing the bilateral LOC reference RDM to the floating RDMs in the ipsilateral conditions. The calculation followed the same procedure used in the decoding and RSA analyses. Specifically, for the left hemisphere, the RSA results are derived from the LHLV sub-condition (left hemisphere ROI, left visual field), and for the right hemisphere, the RSA results are from the RHRV sub-condition (right hemisphere ROI, right visual field). The RSA was calculated at individual level using their individual reference RDM and floating RDMs for each task.

For visualization on the common surface space, the representational similarity values were averaged across all participants in the fsaverage space. For statistical analysis (Figure S11), *r* values were extracted from the LOC for each participant in native surface space, and all values were Fisher Z-transformed. All Fisher Z-transformed correlation values were pooled across participants to form the population distribution. Additionally, participant-level averages were computed and used in statistical analyses.

#### Cross-ROI partial RSA

The cross-ROI partial RSA was performed using decoding-based RDMs calculated as described above. For the VWM tasks, we computed partial correlations between the ipsilateral and contralateral LOC RDMs and all other ipsilateral or contralateral ROI RDM, with the remaining RDMs serving as covariates. Pearson correlation coefficients were used and subsequently Fisher Z-transformed for statistical testing.

### Statistical Analysis

All analyses employed within-subject statistical tests. Given the small sample size, we opted for the Wilcoxon signed-rank test implemented in Pingouin (Vallat, 2018). In the Wilcoxon test, pairs with zero differences were discarded when calculation. Parallel *t*-test are also reported in the Supplementary to show parametric statistics results. In the decoding analysis, we conducted one-tailed tests to determine whether the decoding accuracy in each condition, after subtracting the chance level, was greater than zero, and whether decoding accuracy was higher for cued versus uncued objects. For the ablation decoding, the same statistical tests were applied for each ablation step. For the RSA results, Fisher Z-transformed values were used, and statistical tests directly evaluated whether the Fisher Z value was greater than zero. No multiple comparisons correction was applied to any MVPA statistical results.

## Acknowledgements

We thank MiYoung Kwon and Zitong Lu for their invaluable feedback on manuscript revisions. We thank Chunfang Yan for helping with collecting fMRI data, and Xinyue Peng for assistance in writing the BA task program.

## Author contributions

P.B., J.Y., and W.L. designed the study; W.L. and J.Y. collected the data; W.L. analyzed the data; W.L., J.Y., and P.B. wrote the original draft; P.B., J.Y., and W.L. reviewed and edited the manuscript.

## Supplementary Figures

**Figure S1.**
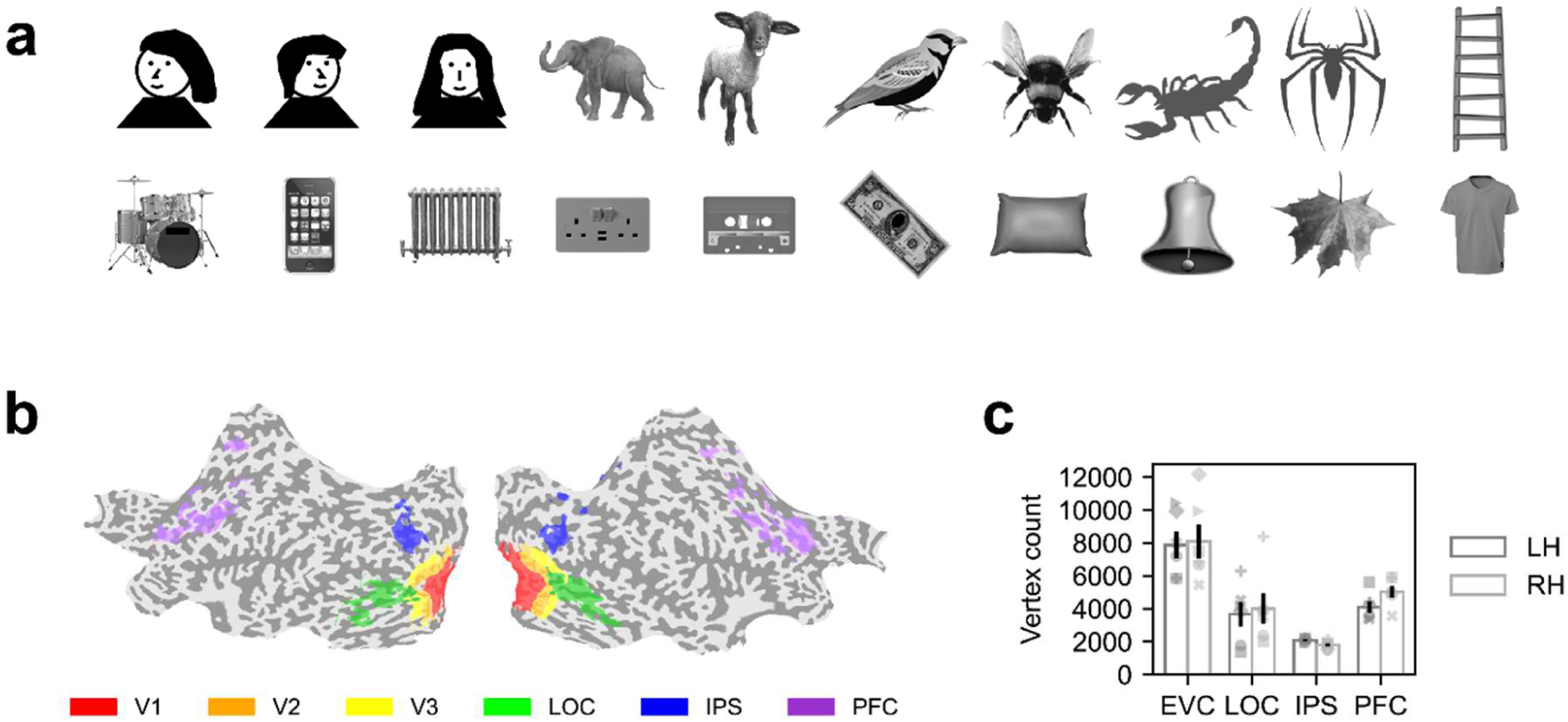
Stimuli and ROI information. Related to Figure 1, 2, 4 and 6. **a.** Twenty grayscale objects used in all experiments. Due to the bioRxiv policy, real face images were replaced with hand-drawn cartoon faces; remaining facial depictions and logos were obscured with black masks. **b.** Cortical ROIs from an example participant. EVC was defined by combining V1–V3. **c.** Average number of vertices per ROI across participants in each hemisphere. Error bars represent ±1 *s.e.m*. Distinct symbols represent individual participants (see Figure S3 for symbol-to-participant correspondence). LH, left hemisphere; RH, right hemisphere; *s.e.m.*, standard error of the mean.

**Figure S2.**
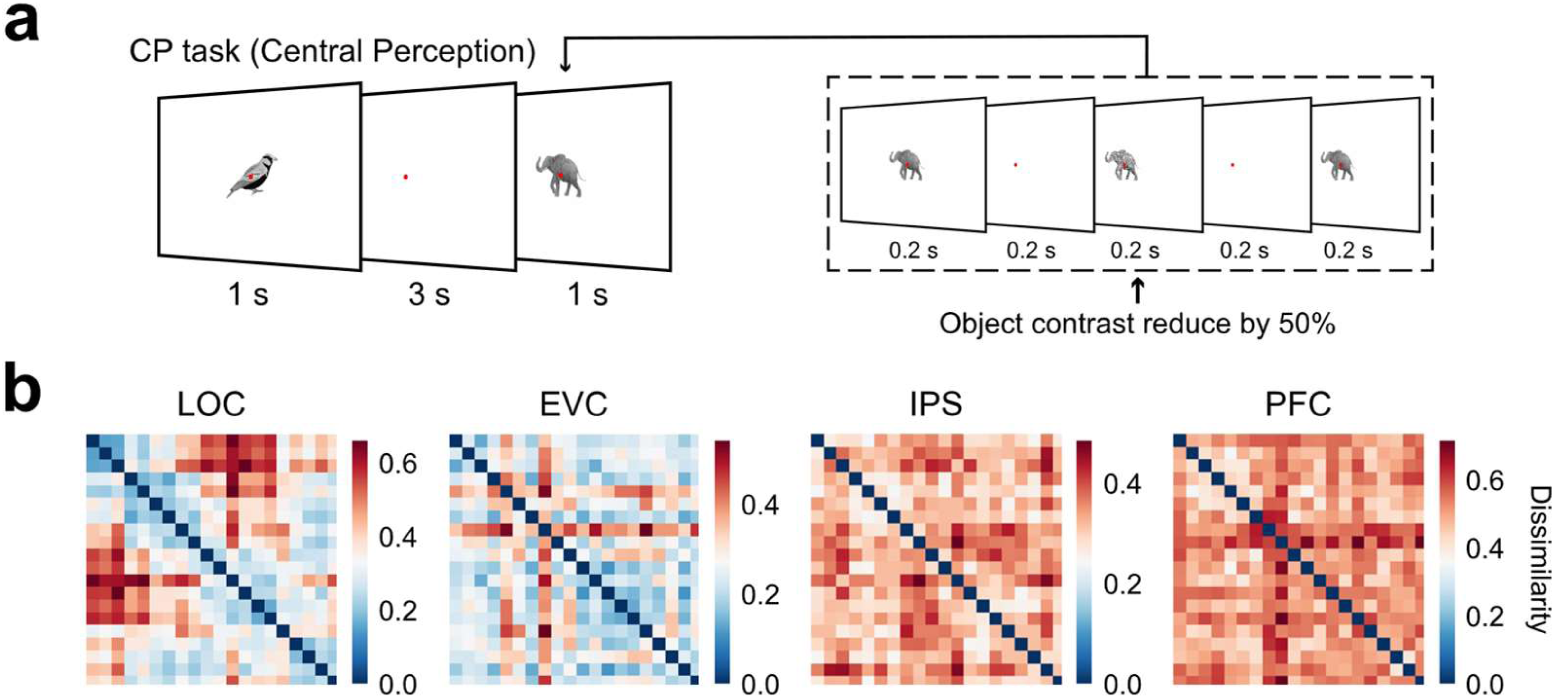
CP task design and RDMs. Related to Figure 1 and 5. **a.** An independent perception (central perception, CP) task designed to serve as a common reference condition for both RSA and searchlight RSA. In this task, an object was presented at the center of the screen. Each trial consisted of three object flashes, and participants were asked to detect a contrast change in the middle frame. **b.** Averaged correlation-based RDMs for each ROI across participants in the CP task. In the RSA analysis, each participant’s individual RDM was used as the independent sensory template.

**Figure S3.**
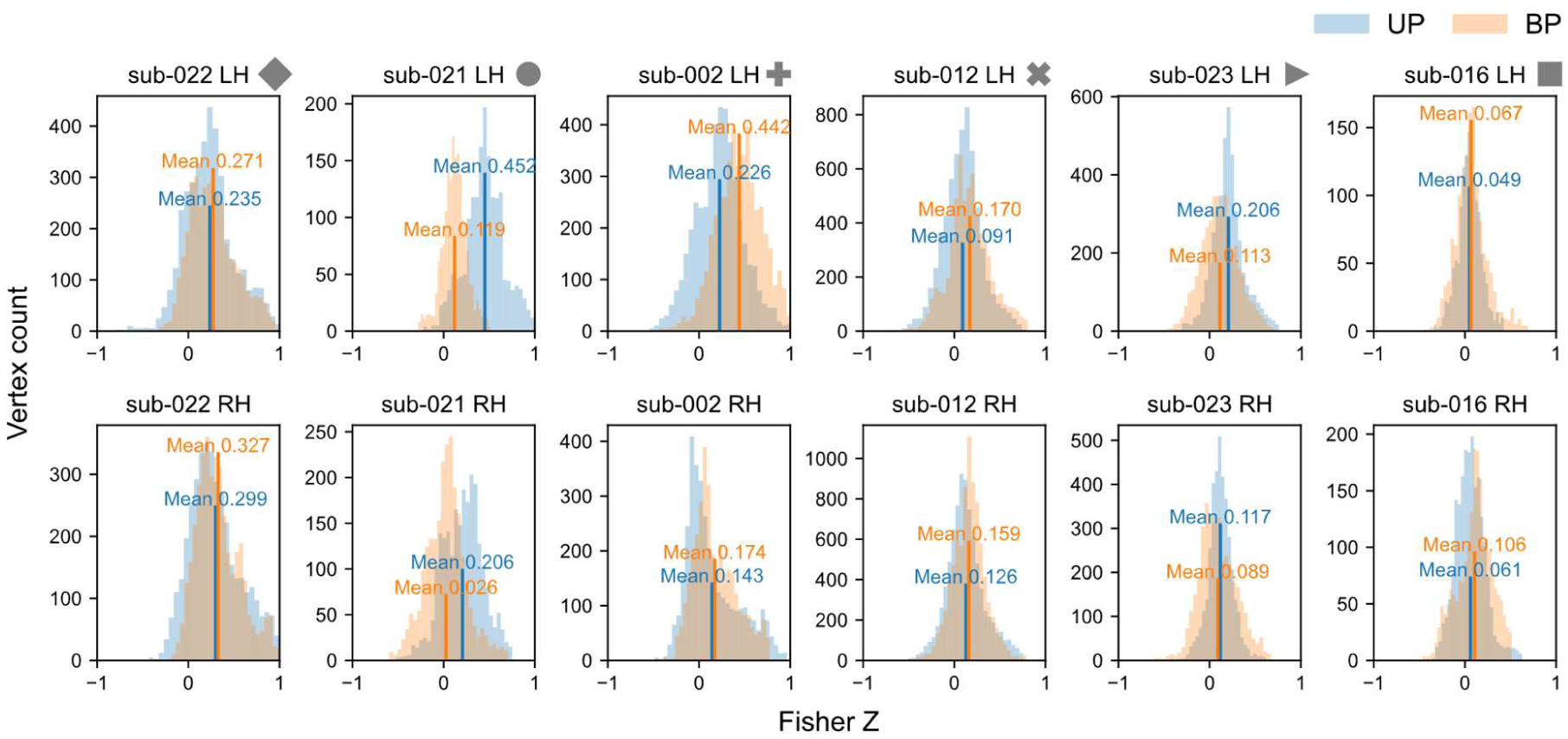
Noise ceiling for UP and BP tasks. Related to Figure 1. The noise ceiling distribution for all vertices within the LOC in UP and BP tasks. The noise ceiling for each vertex was computed by calculating the split-half correlation between trials based on object labels presented in the contralateral visual field. The split-half procedure was repeated 100 times and the results were averaged. Symbols adjacent to subpanel titles denote individual participants, with a consistent symbol-to-participant mapping across all figures. No difference emerged in the averaged noise ceiling between the two tasks across participants (*W* = 10, *p* = 1.000).

**Figure S4.**
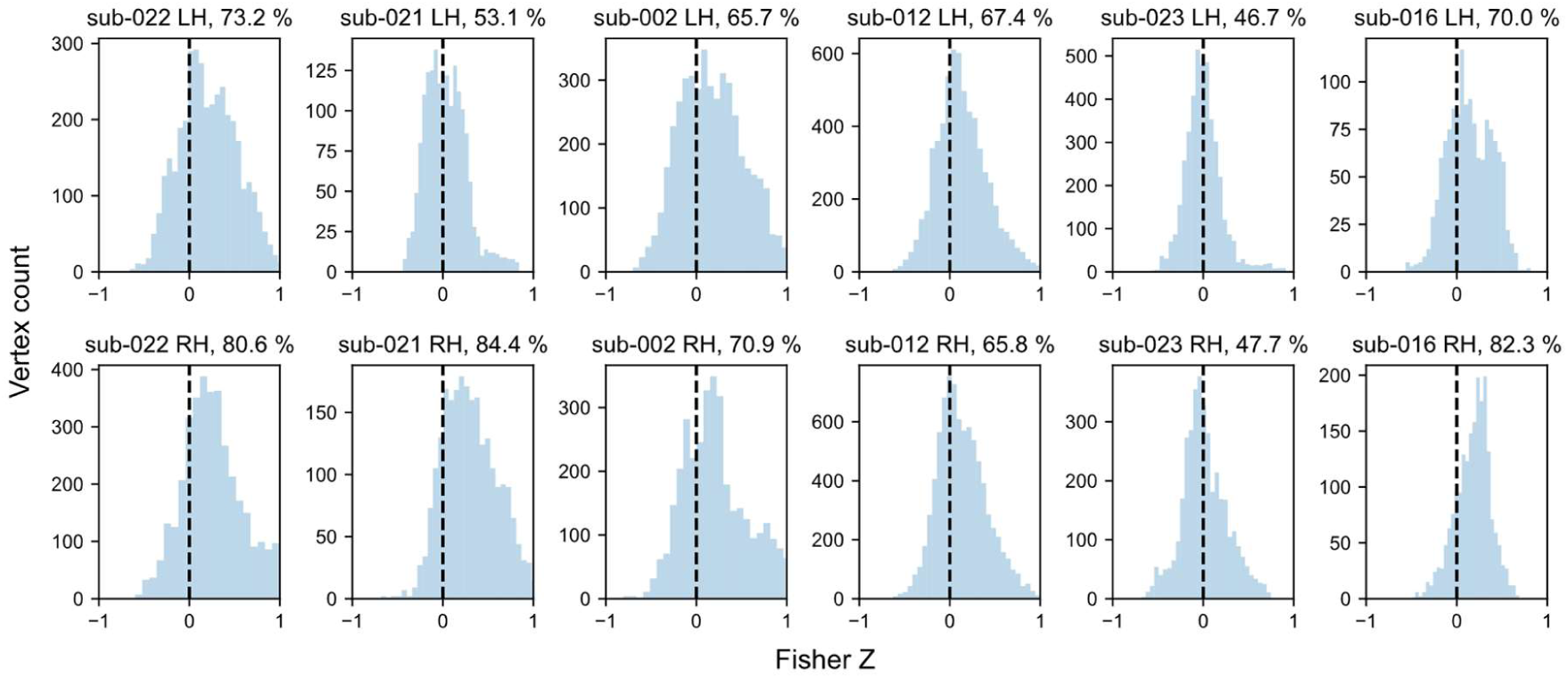
Bilateral responsivity distribution. Related to Figure 3 and 4. Bilateral responsivity distribution in the LOC was calculated from the UP task. For each vertex in the LOC, the bilateral responsivity index was derived by calculating the correlation between the responses to 20 objects in the ipsilateral and contralateral conditions, as described in the Methods. The vertical dashed black lines represent zero, and the percentage in each subpanel title indicates the proportion of vertices with a positive correlation (i.e., the ratio of above-zero vertices).

**Figure S5.**
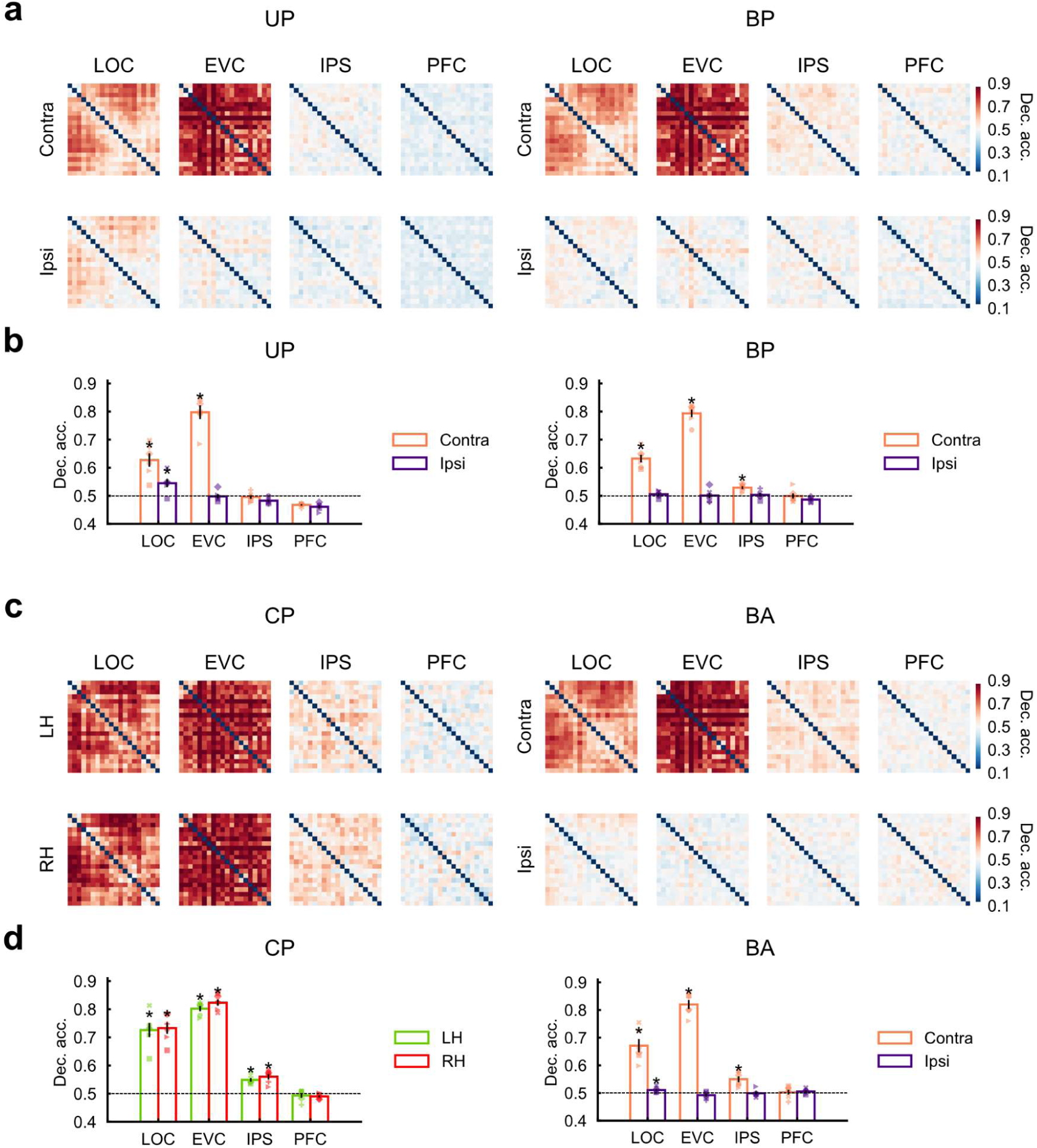
Binary decoding results for perception and attention tasks. Related to Figure 1 and Figure 4. **a.** Decoding-based RDMs for the UP (left) and BP (right) tasks. Each matrix’s value represents pairwise binary decoding accuracy between two of the twenty objects. **b.** Averaged decoding accuracy for the UP (left) and BP (right) tasks across all object pairs, derived from the off-diagonal elements of the RDMs shown in **a**. **c.** Decoding-based RDMs for the CP (left) and BA (right) tasks. Decoding results for the CP task were calculated from the left and right hemisphere data separately, based on the shared central object labels. **d.** Averaged decoding accuracy for the CP (left) and BA (right) tasks. The horizontal dashed black lines indicate the chance-level decoding accuracy. Error bars represent ± 1 *s.e.m*. LH, left hemisphere; RH, right hemisphere. Contra, contralateral; Ipsi, ipsilateral; Dec. acc., decoding accuracy; *s.e.m.*, standard error of the mean. * *p* < 0.05, uncorrected, Wilcoxon signed-rank test.

**Figure S6.**
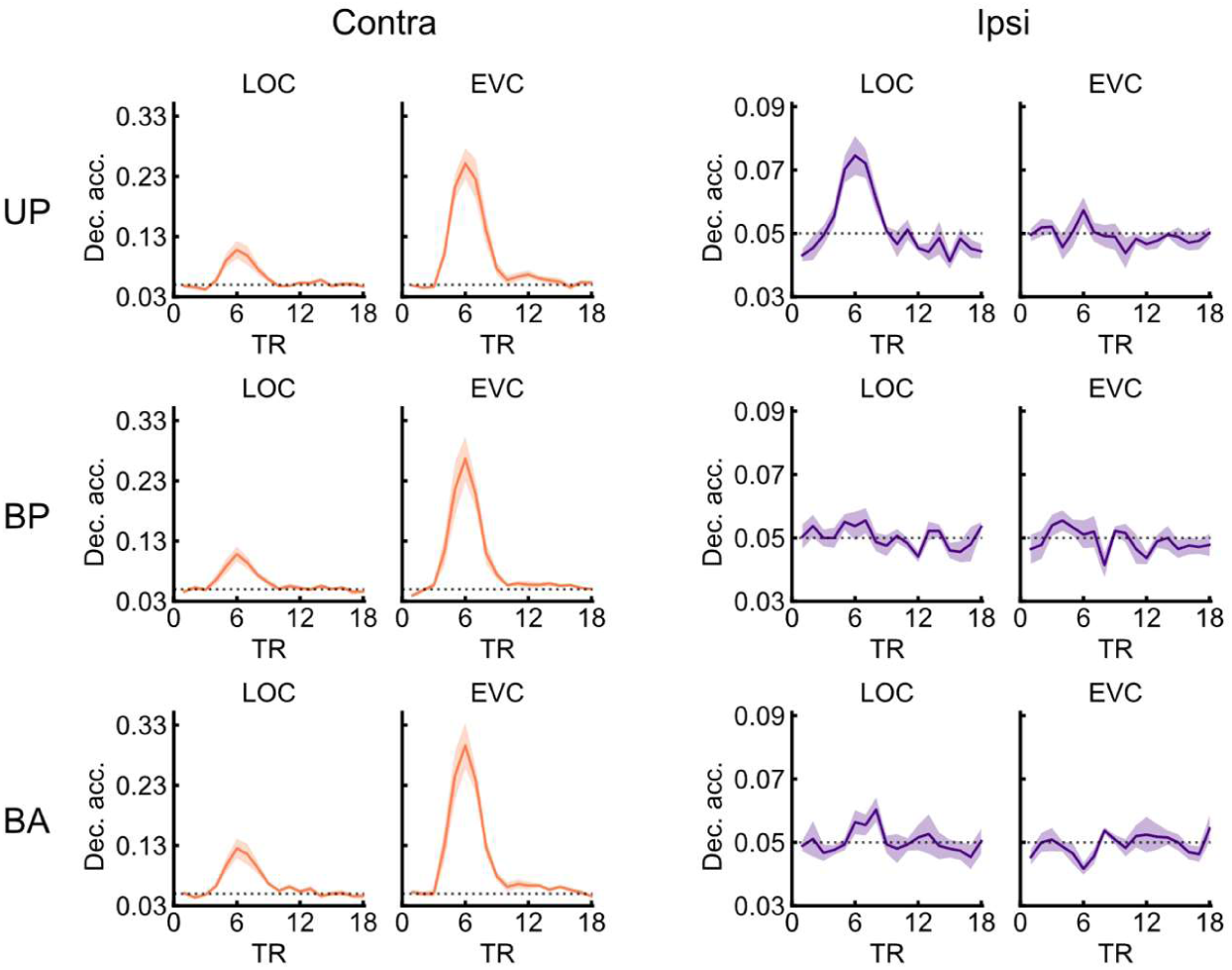
Decoding time course for perception and attention tasks. Related to Figure 1 and Figure 4. We performed a 20-category decoding analysis on the BOLD response for each task. For each trial, we extracted the BOLD signal at stimulus onset and the 17 subsequent TRs, yielding an 18-TR time course per trial similar as in the VWM tasks. For the final two trials of each run, where the full 18-TR window was unavailable, missing TRs were imputed by the mean signal at the corresponding TR across all other trials in that run for each vertex. Shaded areas represent ± 1 *s.e.m*. Contra, contralateral; Ipsi, ipsilateral; Dec. acc., decoding accuracy; *s.e.m.*, standard error of the mean.

**Figure S7.**
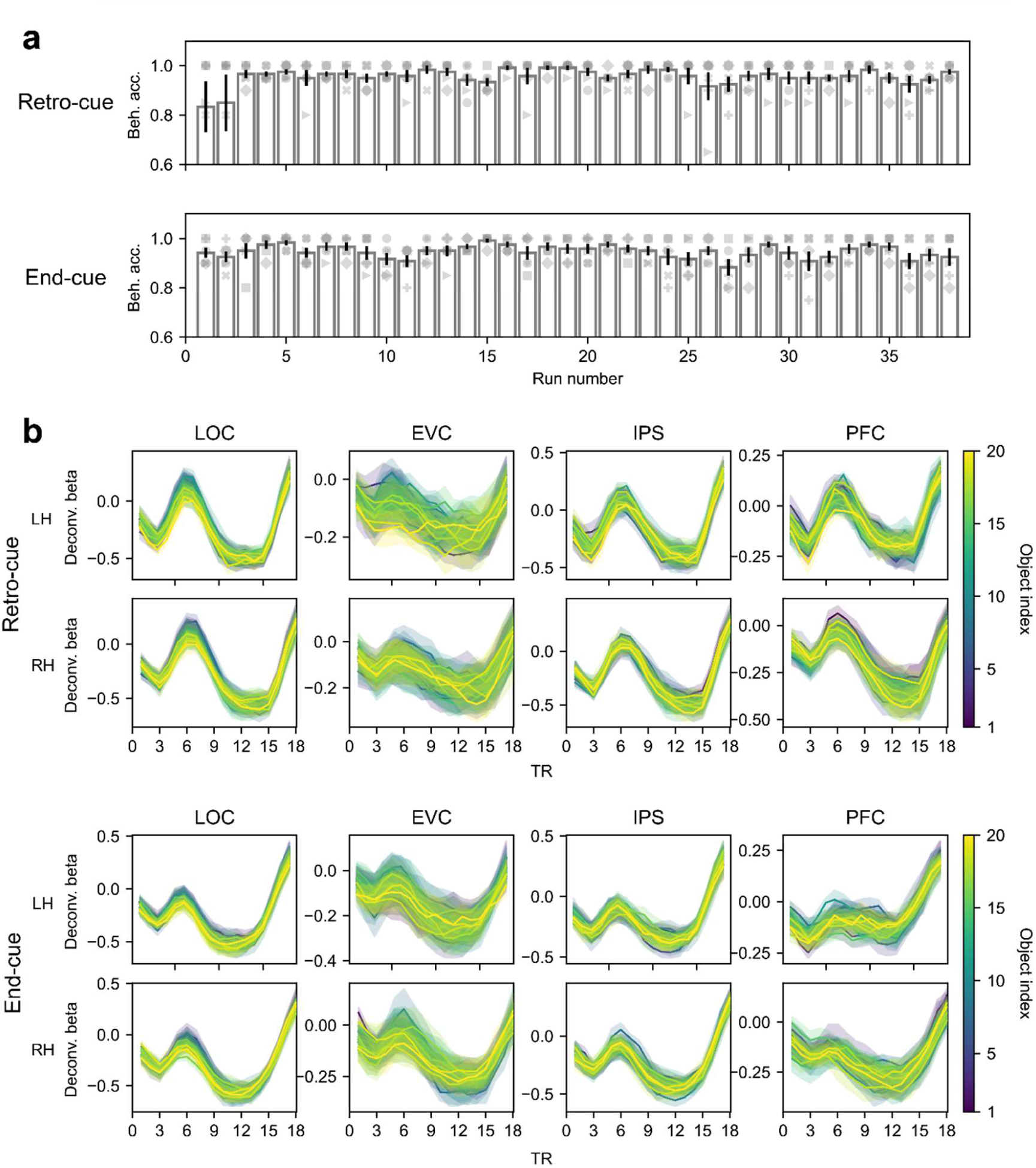
Behavioral performance and deconvolutional time courses in VWM tasks. Related to Figure 2 and 6. **a.** Average behavioral accuracy for each fMRI run in the VWM tasks. **b.** Deconvolutional time courses for VWM tasks. The design matrix used for deconvolution was based on contralateral object labels in the encoding array relative to the hemisphere. Error bars and shaded areas indicate ±1 *s.e.m*. The object index (denoted by color) corresponds to the order shown in Figure S1a, with the top row representing objects 1–10, and the bottom row representing objects 11–20, ordered from left to right. Beh. acc., behavioral accuracy; Deconv. Beta, deconvolved beta; *s.e.m.*, standard error of the mean.

**Figure S8.**
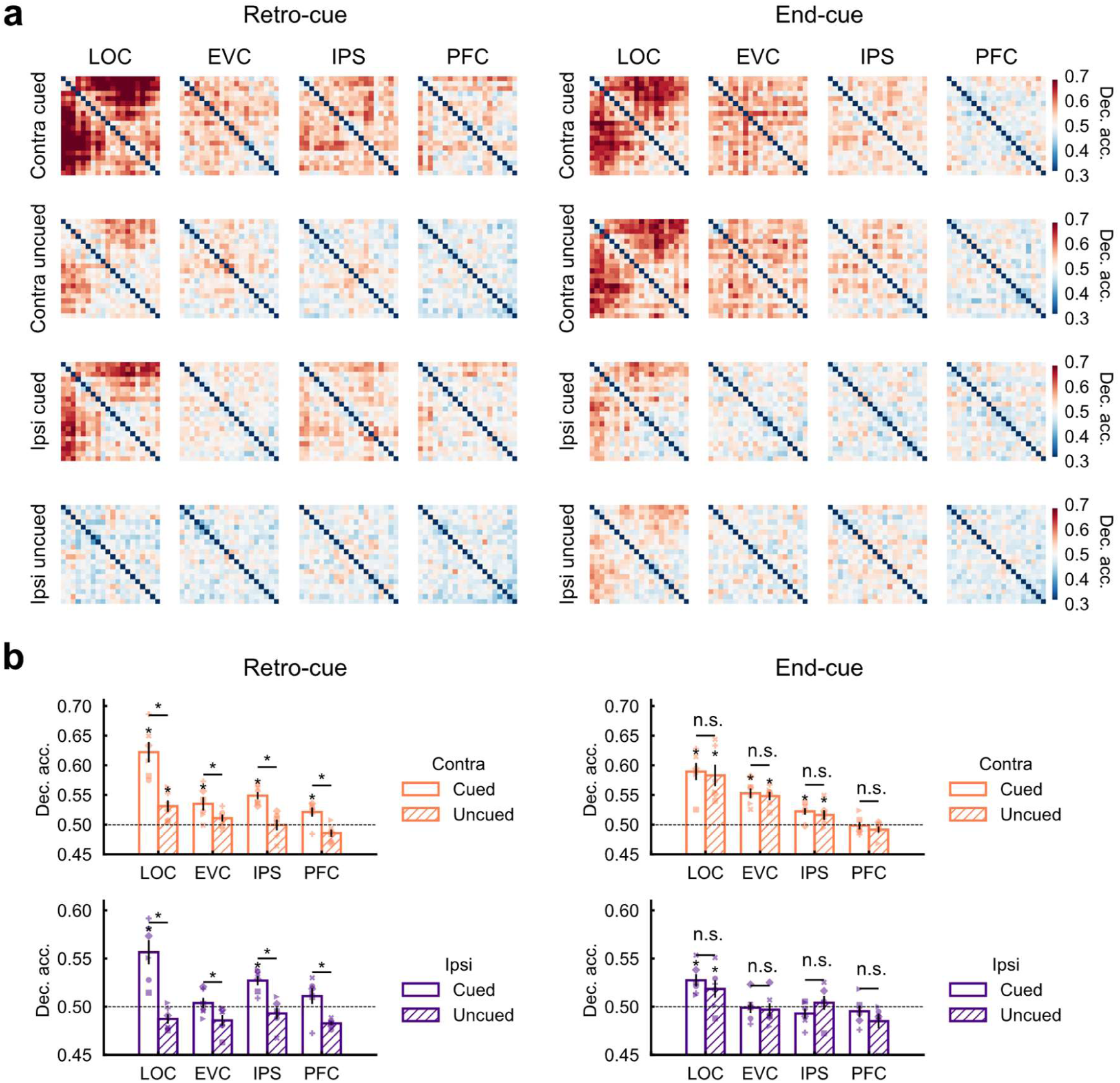
Binary decoding results for VWM tasks. Related to Figure 2 and 6. **a.** Decoding-based RDMs for the retro-cue (left) and end-cue (right) VWM tasks. Each matrix’s value represents pairwise binary decoding accuracy between two of the twenty objects, based on the averaged BOLD response from 6–10 TRs. **b.** Averaged decoding accuracy for the retro-cue (left) and end-cue (right) VWM tasks across all object pairs, derived from the off-diagonal elements of the RDMs shown in **a**. Error bars represent ± 1 *s.e.m*. The horizontal dashed black lines indicate chance level. Contra, contralateral; Ipsi, ipsilateral; Dec. acc., decoding accuracy; *s.e.m.*, standard error of the mean.

**Figure S9.**
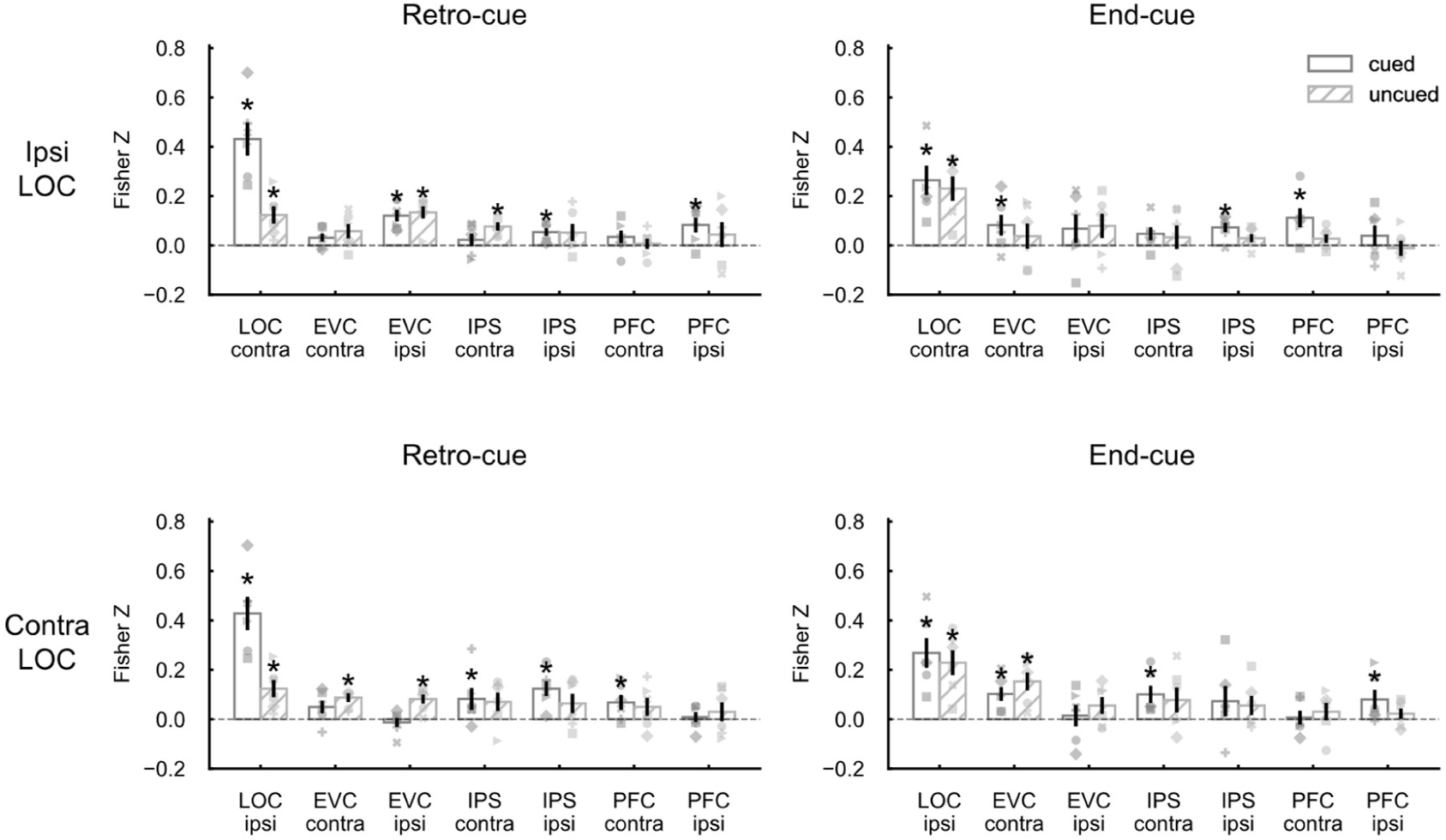
Cross-region partial RSA results for VWM tasks. Related to Figure 6. Cross-ROI partial RSA results for the retro-cue and end-cue VWM tasks. **a.** Partial RSA results between the ipsilateral LOC and other regions. **b.** Partial RSA results between the contralateral LOC and other regions. All RDMs were derived from decoding-based measures as shown in **Figure S8a**. RSA correlation values were Fisher Z-transformed. Error bars represent ± 1 *s.e.m*. Contra, contralateral; Ipsi, ipsilateral; *s.e.m.*, standard error of the mean. * *p* < 0.05, uncorrected, comparing each condition to zero, Wilcoxon signed-rank test.

**Figure S10.**
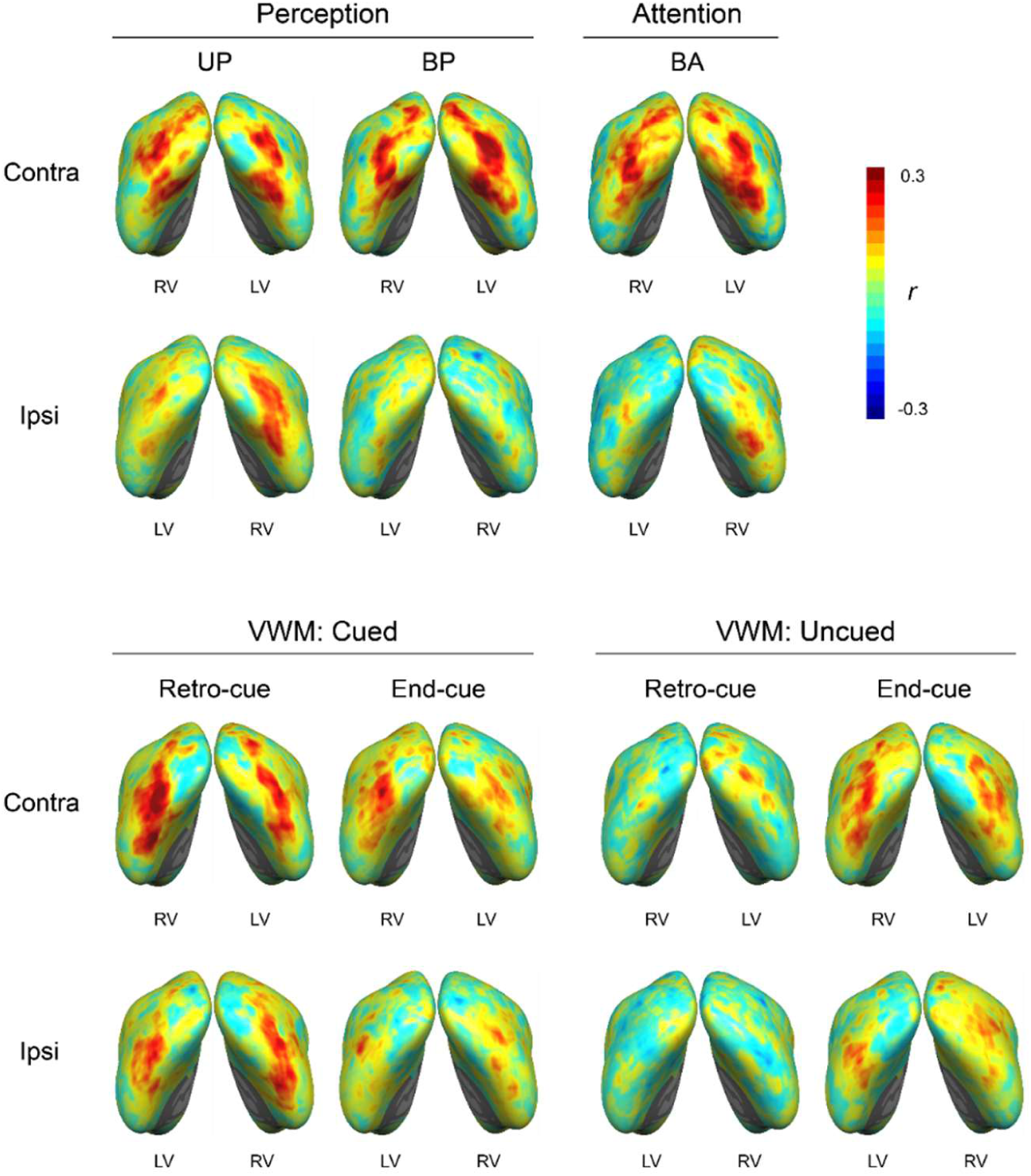
Searchlight RSA results in the fsaverage space. Related to Figure 5. Similar to Figure 5, but displaying both contralateral and ipsilateral results without truncation. Contra, contralateral; Ipsi, ipsilateral; LV, left visual field; RV, right visual field.

**Figure S11.**
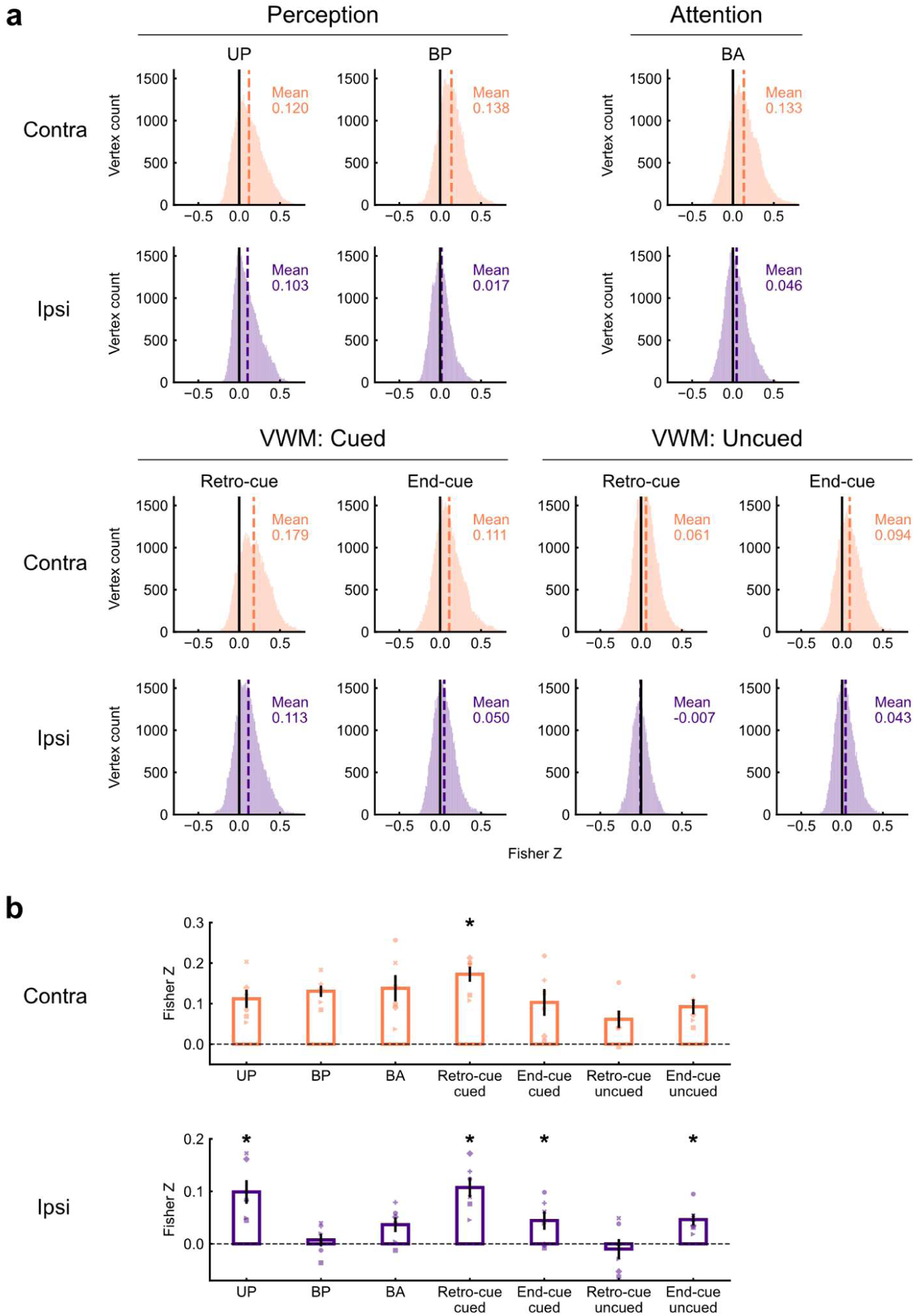
Searchlight RSA results in participants’ native surface space. Related to Figure 5. **a.** Searchlight RSA correlation values distribution within LOC summed across all participants in their native surface space and ROI. The number in each subpanel is the averaged correlation value across all vertices for all participants. The vertical black lines represent zero, while the vertical dashed colored lines represent the mean correlation. **b.** Averaged correlation values within LOC across participants in different task conditions. Error bars represent ± 1 *s.e.m.* Contra, contralateral; Ipsi, ipsilateral; *s.e.m.*, standard error of the mean. * *p* < 0.05, uncorrected, representing whether the correlation values in the current task condition are significantly higher than the BP task.

## Supplementary Tables

**Table S1.** Statistical results for decoding accuracy in the perception tasks. Related to Figure 1.

| Task | ROI | Condition | $M \pm s.e.m$ | $W$ | $W-p$ | $W-RBC$ | $W-CLES$ | $t$ | $t-p$ |
| --- | --- | --- | --- | --- | --- | --- | --- | --- | --- |
| UP | LOC | Contra | $0.105 \pm 0.015$ | **21\*** | **0.016** | 1.000 | 1.000 | **3.615\*\*** | **0.008** |
| UP | LOC | Ipsi | $0.066 \pm 0.006$ | **19\*** | **0.047** | 0.810 | 0.833 | **2.533\*** | **0.026** |
| UP | EVC | Contra | $0.300 \pm 0.032$ | **21\*** | **0.016** | 1.000 | 1.000 | **7.858\*\*\*** | **0.000** |
| UP | EVC | Ipsi | $0.045 \pm 0.005$ | 5.5 | 0.891 | -0.476 | 0.333 | -0.990 | 0.816 |
| BP | LOC | Contra | $0.116 \pm 0.015$ | **21\*** | **0.016** | 1.000 | 1.000 | **4.506\*\*** | **0.003** |
| BP | LOC | Ipsi | $0.053 \pm 0.002$ | 18 | 0.078 | 0.714 | 0.833 | 1.908 | 0.057 |
| BP | EVC | Contra | $0.277 \pm 0.018$ | **21\*** | **0.016** | 1.000 | 1.000 | **12.946\*\*\*** | **0.000** |
| BP | EVC | Ipsi | $0.054 \pm 0.006$ | 11 | 0.500 | 0.048 | 0.333 | 0.688 | 0.261 |

*Note.* One-sample tests comparing decoding accuracy against chance level. Contra, contralateral; Ipsi, ipsilateral. $M \pm s.e.m.$ , mean ± standard error of the mean. $W$ , Wilcoxon signed-rank statistic; $W-RBC$ , rank-biserial correlation effect size; $W-CLES$ , common language effect size. $t$ , t-statistic. \* $p < 0.05$ , \*\* $p < 0.01$ , \*\*\* $p < 0.001$ . Statistic values reaching significance ( $p < 0.05$ ) are shown in boldface.

**Table S2.** Statistical results for RSA in the perception tasks. Related to Figure 1.

| task | ROI | Condition | $M \pm s.e.m$ | $W$ | $W-p$ | $W-RBC$ | $W-CLES$ | $t$ | $t-p$ |
| --- | --- | --- | --- | --- | --- | --- | --- | --- | --- |
| UP | LOC | Contra | $0.360 \pm 0.082$ | **21\*** | **0.016** | 1.000 | 1.000 | **4.376\*\*\*** | **0.004** |
| UP | LOC | Ipsi | $0.328 \pm 0.091$ | **21\*** | **0.016** | 1.000 | 1.000 | **3.610\*\*** | **0.008** |
| UP | EVC | Contra | $0.618 \pm 0.020$ | **21\*** | **0.016** | 1.000 | 1.000 | **31.044\*\*\*** | **0.000** |
| UP | EVC | Ipsi | $0.039 \pm 0.022$ | 18 | 0.078 | 0.714 | 0.833 | 1.761 | 0.069 |
| BP | LOC | Contra | $0.367 \pm 0.056$ | **21\*** | **0.016** | 1.000 | 1.000 | **6.542\*\*\*** | **0.001** |
| BP | LOC | Ipsi | $0.035 \pm 0.038$ | 14 | 0.281 | 0.333 | 0.500 | 0.907 | 0.203 |
| BP | EVC | Contra | $0.676 \pm 0.041$ | **21\*** | **0.016** | 1.000 | 1.000 | **16.676\*\*\*** | **0.000** |
| BP | EVC | Ipsi | $0.082 \pm 0.056$ | 16 | 0.156 | 0.524 | 0.667 | 1.468 | 0.101 |

*Note.* One-sample tests comparing the mean Fisher Z-transformed correlation coefficient against zero. Contra, contralateral; Ipsi, ipsilateral. $M \pm s.e.m.$ , mean ± standard error of the mean. $W$ , Wilcoxon signed-rank statistic; $W-RBC$ , rank-biserial correlation effect size; $W-CLES$ , common language effect size. $t$ , t-statistic. \* $p < 0.05$ , \*\* $p < 0.01$ , \*\*\* $p < 0.001$ . Statistic values reaching significance ( $p < 0.05$ ) are shown in boldface.

**Table S3.** Statistical results for decoding accuracy in the retro-cue VWM task. Related to Figure 2 and 6.

| ROI | Condition Lateral | Condition Cue | $M \pm s.e.m$ | $W$ | $W-p$ | $W-RBC$ | $W-CLES$ | $t$ | $t-p$ |
| --- | --- | --- | --- | --- | --- | --- | --- | --- | --- |
| LOC | Contra | Cued | 0.111 ± 0.013 | **21\*** | **0.016** | 1.000 | 1.000 | **4.771\*\*** | **0.003** |
| LOC | Contra | Uncued | 0.063 ± 0.004 | **15\*** | **0.030** | 1.000 | 0.917 | **3.437\*\*** | **0.009** |
| LOC | Ipsi | Cued | 0.077 ± 0.008 | **20\*** | **0.031** | 0.905 | 0.833 | **3.651\*\*** | **0.007** |
| LOC | Ipsi | Uncued | 0.050 ± 0.003 | 8 | 0.719 | -0.238 | 0.333 | -0.000 | 0.500 |
| EVC | Contra | Cued | 0.070 ± 0.006 | **21\*** | **0.016** | 1.000 | 1.000 | **3.316\*** | **0.011** |
| EVC | Contra | Uncued | 0.057 ± 0.003 | **19\*** | **0.047** | 0.810 | 0.833 | **2.431\*** | **0.030** |
| EVC | Ipsi | Cued | 0.049 ± 0.003 | 8 | 0.719 | -0.238 | 0.500 | -0.326 | 0.621 |
| EVC | Ipsi | Uncued | 0.049 ± 0.004 | 9 | 0.656 | -0.143 | 0.500 | -0.151 | 0.557 |
| IPS | Contra | Cued | 0.071 ± 0.003 | **21\*** | **0.016** | 1.000 | 1.000 | **7.665\*\*\*** | **0.000** |
| IPS | Contra | Uncued | 0.052 ± 0.003 | 10 | 0.295 | 0.333 | 0.750 | 0.455 | 0.334 |
| IPS | Ipsi | Cued | 0.063 ± 0.003 | **20\*** | **0.031** | 0.905 | 0.833 | **4.333\*\*** | **0.004** |
| IPS | Ipsi | Uncued | 0.050 ± 0.001 | 9 | 0.656 | -0.143 | 0.500 | 0.299 | 0.389 |
| PFC | Contra | Cued | 0.059 ± 0.004 | **19\*** | **0.047** | 0.810 | 0.833 | **2.263\*** | **0.037** |
| PFC | Contra | Uncued | 0.046 ± 0.005 | 4 | 0.922 | -0.619 | 0.167 | -0.876 | 0.789 |
| PFC | Ipsi | Cued | 0.063 ± 0.004 | **21\*** | **0.016** | 1.000 | 1.000 | **3.496\*\*** | **0.009** |
| PFC | Ipsi | Uncued | 0.047 ± 0.002 | 4 | 0.922 | -0.619 | 0.167 | -1.481 | 0.901 |

*Note.* One-sample tests comparing decoding accuracy against chance level. Contra, contralateral; Ipsi, ipsilateral. $M \pm s.e.m.$ , mean ± standard error of the mean. $W$ , Wilcoxon signed-rank statistic; $W-RBC$ , rank-biserial correlation effect size; $W-CLES$ , common language effect size. $t$ , t-statistic. \* $p < 0.05$ , \*\* $p < 0.01$ , \*\*\* $p < 0.001$ . Statistic values reaching significance ( $p < 0.05$ ) are shown in boldface.

**Table S4.** Statistical results for decoding accuracy in the retro-cue VWM task. Related to Figure 2 and 6.

| ROI | Condition | *W* | *W-p* | *W-RBC* | *W-CLES* | *t* | *t-p* |
| --- | --- | --- | --- | --- | --- | --- | --- |
| LOC | Contra | **21\*** | **0.016** | 1.000 | 1.000 | **4.870\*\*** | **0.002** |
| LOC | Ipsi | **19\*** | **0.047** | 0.810 | 0.889 | **2.854\*** | **0.018** |
| EVC | Contra | 18 | 0.078 | 0.714 | 0.681 | 1.831 | 0.063 |
| EVC | Ipsi | 10 | 0.578 | -0.048 | 0.458 | -0.035 | 0.513 |
| IPS | Contra | **21\*** | **0.016** | 1.000 | 1.000 | **3.568\*\*** | **0.008** |
| IPS | Ipsi | **21\*** | **0.016** | 1.000 | 0.861 | **5.534\*\*** | **0.001** |
| PFC | Contra | **20\*** | **0.031** | 0.905 | 0.778 | **2.832\*** | **0.018** |
| PFC | Ipsi | **21\*** | **0.016** | 1.000 | 0.972 | **4.142\*\*** | **0.004** |

*Note.* Paired-sample tests comparing decoding accuracy in the cued against uncued condition. Contra, contralateral; Ipsi, ipsilateral. *W*, Wilcoxon signed-rank statistic; *W-RBC*, rank-biserial correlation effect size; *W-CLES*, common language effect size. *t*, t-statistic. \* $p < 0.05$ , \*\* $p < 0.01$ . Statistic values reaching significance ( $p < 0.05$ ) are shown in boldface.

**Table S5.** Statistical results for decoding accuracy in the end-cue VWM task. Related to Figure 2 and 6.

| ROI | Condition Lateral | Condition Cue | $M \pm s.e.m$ | $W$ | $W-p$ | $W-RBC$ | $W-CLES$ | $t$ | $t-p$ |
| --- | --- | --- | --- | --- | --- | --- | --- | --- | --- |
| LOC | Contra | Cued | 0.087 ± 0.007 | **21\*** | **0.016** | 1.000 | 1.000 | **5.062\*\*** | **0.002** |
| LOC | Contra | Uncued | 0.085 ± 0.007 | **21\*** | **0.016** | 1.000 | 1.000 | **4.783\*\*** | **0.002** |
| LOC | Ipsi | Cued | 0.066 ± 0.004 | **21\*** | **0.016** | 1.000 | 1.000 | **4.568\*\*** | **0.003** |
| LOC | Ipsi | Uncued | 0.063 ± 0.004 | **21\*** | **0.016** | 1.000 | 1.000 | **3.612\*\*** | **0.008** |
| EVC | Contra | Cued | 0.073 ± 0.006 | **21\*** | **0.016** | 1.000 | 1.000 | **3.817\*\*** | **0.006** |
| EVC | Contra | Uncued | 0.074 ± 0.006 | **21\*** | **0.016** | 1.000 | 1.000 | **4.104\*\*** | **0.005** |
| EVC | Ipsi | Cued | 0.051 ± 0.004 | 10 | 0.578 | -0.048 | 0.500 | 0.267 | 0.400 |
| EVC | Ipsi | Uncued | 0.048 ± 0.003 | 7 | 0.781 | -0.333 | 0.333 | -0.681 | 0.737 |
| IPS | Contra | Cued | 0.062 ± 0.003 | **20\*** | **0.031** | 0.905 | 0.833 | **4.330\*\*** | **0.004** |
| IPS | Contra | Uncued | 0.065 ± 0.007 | **19\*** | **0.047** | 0.810 | 0.833 | **2.032\*** | **0.049** |
| IPS | Ipsi | Cued | 0.050 ± 0.003 | 9 | 0.656 | -0.143 | 0.333 | -0.000 | 0.500 |
| IPS | Ipsi | Uncued | 0.054 ± 0.003 | 17 | 0.109 | 0.619 | 0.667 | 1.663 | 0.079 |
| PFC | Contra | Cued | 0.052 ± 0.002 | 13 | 0.344 | 0.238 | 0.500 | 0.794 | 0.232 |
| PFC | Contra | Uncued | 0.052 ± 0.004 | 13 | 0.344 | 0.238 | 0.500 | 0.611 | 0.284 |
| PFC | Ipsi | Cued | 0.051 ± 0.002 | 11 | 0.500 | 0.048 | 0.500 | 0.374 | 0.362 |
| PFC | Ipsi | Uncued | 0.049 ± 0.003 | 7 | 0.781 | -0.333 | 0.333 | -0.341 | 0.627 |

*Note.* One-sample tests comparing decoding accuracy against chance level. Contra, contralateral; Ipsi, ipsilateral. $M \pm s.e.m.$ , mean ± standard error of the mean. $W$ , Wilcoxon signed-rank statistic; $W-RBC$ , rank-biserial correlation effect size; $W-CLES$ , common language effect size. $t$ , t-statistic. \* $p < 0.05$ , \*\* $p < 0.01$ . Statistic values reaching significance ( $p < 0.05$ ) are shown in boldface.

**Table S6.**
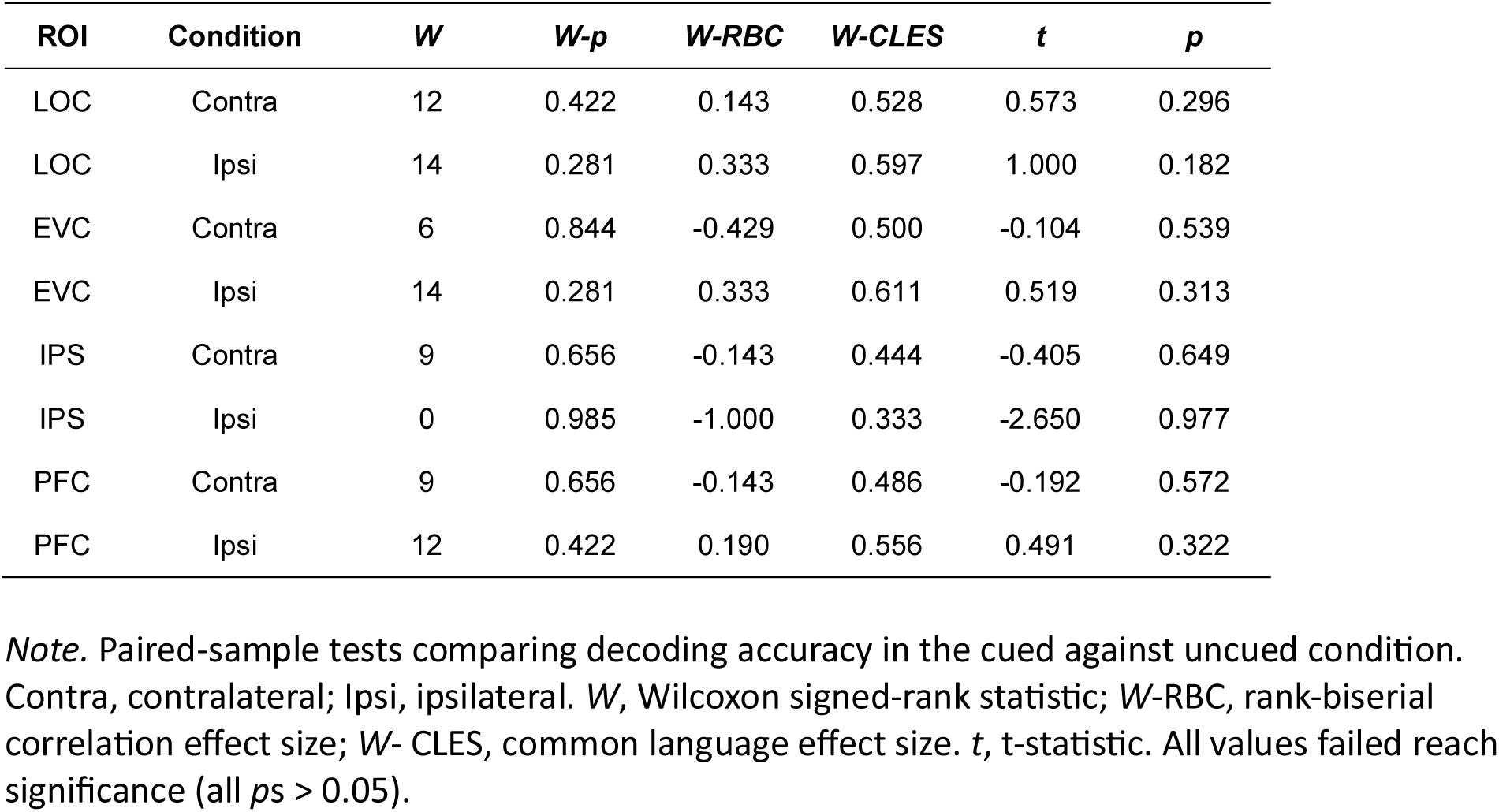
Statistical results for decoding accuracy in the end-cue VWM task. Related to Figure 2 and 6.

**Table S7.** Statistical results for decoding accuracy in the Bilateral Attention (BA) task. Related to Figure 4.

| ROI | Condition | $M \pm s.e.m$ | $W$ | $W-p$ | $W-RBC$ | $W-CLES$ | $t$ | $p$ |
| --- | --- | --- | --- | --- | --- | --- | --- | --- |
| LOC | Contra | $0.139 \pm 0.020$ | **21\*** | **0.016** | 1.000 | 1.000 | **4.380\*\*** | **0.004** |
| LOC | Ipsi | $0.055 \pm 0.002$ | **20\*** | **0.031** | 0.905 | 0.833 | **2.421\*** | **0.030** |
| EVC | Contra | $0.338 \pm 0.026$ | **21\*** | **0.016** | 1.000 | 1.000 | **10.870\*\*\*** | **0.000** |
| EVC | Ipsi | $0.044 \pm 0.003$ | 3 | 0.953 | -0.714 | 0.333 | -1.537 | 0.908 |

*Note.* One-sample tests comparing decoding accuracy against chance level. Contra, contralateral; Ipsi, ipsilateral. $M \pm s.e.m.$ , mean ± standard error of the mean. $W$ , Wilcoxon signed-rank statistic; $W-RBC$ , rank-biserial correlation effect size; $W-CLES$ , common language effect size. $t$ , t-statistic. \* $p < 0.05$ , \*\* $p < 0.01$ , \*\*\* $p < 0.001$ . Statistic values reaching significance ( $p < 0.05$ ) are shown in boldface.

**Table S8.**
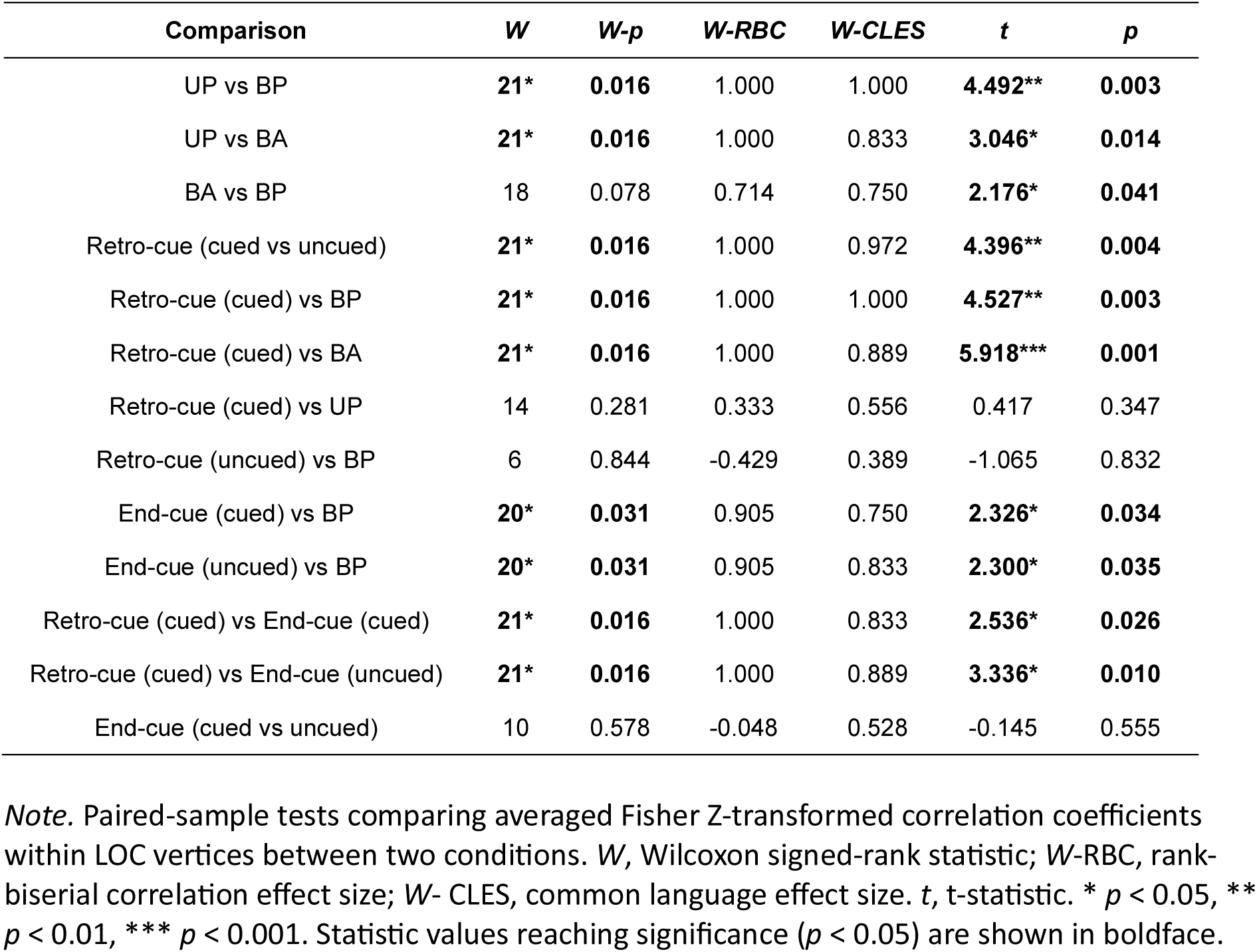
Statistical results for sensory-based ipsilateral representation strength from searchlight RSA. Related to Figure 5.

| Comparison | *W* | *W-p* | *W-RBC* | *W-CLES* | *t* | *p* |
| --- | --- | --- | --- | --- | --- | --- |
| UP vs BP | **21\*** | **0.016** | 1.000 | 1.000 | **4.492\*\*** | **0.003** |
| UP vs BA | **21\*** | **0.016** | 1.000 | 0.833 | **3.046\*** | **0.014** |
| BA vs BP | 18 | 0.078 | 0.714 | 0.750 | **2.176\*** | **0.041** |
| Retro-cue (cued vs uncued) | **21\*** | **0.016** | 1.000 | 0.972 | **4.396\*\*** | **0.004** |
| Retro-cue (cued) vs BP | **21\*** | **0.016** | 1.000 | 1.000 | **4.527\*\*** | **0.003** |
| Retro-cue (cued) vs BA | **21\*** | **0.016** | 1.000 | 0.889 | **5.918\*\*\*** | **0.001** |
| Retro-cue (cued) vs UP | 14 | 0.281 | 0.333 | 0.556 | 0.417 | 0.347 |
| Retro-cue (uncued) vs BP | 6 | 0.844 | -0.429 | 0.389 | -1.065 | 0.832 |
| End-cue (cued) vs BP | **20\*** | **0.031** | 0.905 | 0.750 | **2.326\*** | **0.034** |
| End-cue (uncued) vs BP | **20\*** | **0.031** | 0.905 | 0.833 | **2.300\*** | **0.035** |
| Retro-cue (cued) vs End-cue (cued) | **21\*** | **0.016** | 1.000 | 0.833 | **2.536\*** | **0.026** |
| Retro-cue (cued) vs End-cue (uncued) | **21\*** | **0.016** | 1.000 | 0.889 | **3.336\*** | **0.010** |
| End-cue (cued vs uncued) | 10 | 0.578 | -0.048 | 0.528 | -0.145 | 0.555 |

*Note.* Paired-sample tests comparing averaged Fisher Z-transformed correlation coefficients within LOC vertices between two conditions. *W*, Wilcoxon signed-rank statistic; *W-RBC*, rank-biserial correlation effect size; *W-CLES*, common language effect size. *t*, t-statistic. \* $p < 0.05$ , \*\* $p < 0.01$ , \*\*\* $p < 0.001$ . Statistic values reaching significance ( $p < 0.05$ ) are shown in boldface.

## Reference

Bao, P., She, L., McGill, M., & Tsao, D. Y. (2020). A map of object space in primate inferotemporal cortex. Nature, 583(7814), Article 7814. 10.1038/s41586-020-2350-5

Bao, P., & Tsao, D. Y. (2018). Representation of multiple objects in macaque category-selective areas. Nature Communications, 9(1), Article 1. 10.1038/s41467-018-04126-7

Benson, N. C., Jamison, K. W., & Arcaro, M. J. (2018). The Human Connectome Project 7 Tesla retinotopy dataset: Description and population receptive field analysis. Journal of Vision, 22. 10.1167/18.13.23

Bettencourt, K. C., & Xu, Y. (2016). Decoding the content of visual short-term memory under distraction in occipital and parietal areas. Nature Neuroscience, 19(1), Article 1. 10.1038/nn.4174

Brainard, D. H. (1997). The Psychophysics Toolbox. Spatial Vision, 10(4), 433–436. 10.1163/156856897X00357

Brincat, S. L., Donoghue, J. A., Mahnke, M. K., Kornblith, S., Lundqvist, M., & Miller, E. K. (2021). Interhemispheric transfer of working memories. Neuron, 109(6), 1055–1066.e4. 10.1016/j.neuron.2021.01.016

Christophel, T. B., Klink, P. C., Spitzer, B., Roelfsema, P. R., & Haynes, J.-D. (2017). The Distributed Nature of Working Memory. Trends in Cognitive Sciences, 21(2), 111–124. 10.1016/j.tics.2016.12.007

Cichy, R. M., Chen, Y., & Haynes, J.-D. (2011). Encoding the identity and location of objects in human LOC. NeuroImage, 54(3), 2297–2307. 10.1016/j.neuroimage.2010.09.044

Cichy, R. M., Sterzer, P., Heinzle, J., Elliott, L. T., Ramirez, F., & Haynes, J.-D. (2013). Probing principles of large-scale object representation: Category preference and location encoding. Human Brain Mapping, 34(7), 1636–1651. 10.1002/hbm.22020

D’Esposito, M., & Postle, B. R. (2015). The Cognitive Neuroscience of Working Memory. Annual Review of Psychology, 66(1), 115–142. 10.1146/annurev-psych-010814-015031

DeYoe, E. A., Carman, G. J., Bandettini, P., Glickman, S., Wieser, J., Cox, R., Miller, D., & Neitz, J. (1996). Mapping striate and extrastriate visual areas in human cerebral cortex. Proceedings of the National Academy of Sciences, 93(6), 2382–2386. 10.1073/pnas.93.6.2382

Emrich, S. M., Riggall, A. C., LaRocque, J. J., & Postle, B. R. (2013). Distributed Patterns of Activity in Sensory Cortex Reflect the Precision of Multiple Items Maintained in Visual Short-Term Memory. Journal of Neuroscience, 33(15), 6516–6523. 10.1523/JNEUROSCI.5732-12.2013

Engel, S. A., Glover, G. H., & Wandell, B. A. (1997). Retinotopic organization in human visual cortex and the spatial precision of functional MRI. Cerebral Cortex, 7(2), 181–192. 10.1093/cercor/7.2.181

Esteban, O., Markiewicz, C. J., Blair, R. W., Moodie, C. A., Isik, A. I., Erramuzpe, A., Kent, J. D., Goncalves, M., DuPre, E., Snyder, M., Oya, H., Ghosh, S. S., Wright, J., Durnez, J., Poldrack, R. A., & Gorgolewski, K. J. (2019). fMRIPrep: A robust preprocessing pipeline for functional MRI. Nature Methods, 16(1), 111–116. 10.1038/s41592-018-0235-4

Ester, E. F., Serences, J. T., & Awh, E. (2009). Spatially Global Representations in Human Primary Visual Cortex during Working Memory Maintenance. Journal of Neuroscience, 29(48), 15258–15265. 10.1523/JNEUROSCI.4388-09.2009

Fukuda, K., Kang, M.-S., & Woodman, G. F. (2016). Distinct neural mechanisms for spatially lateralized and spatially global visual working memory representations. Journal of Neurophysiology, 116(4), 1715–1727. 10.1152/jn.00991.2015

Glasser, M. F., Coalson, T. S., Robinson, E. C., Hacker, C. D., Harwell, J., Yacoub, E., Ugurbil, K., Andersson, J., Beckmann, C. F., Jenkinson, M., Smith, S. M., & Van Essen, D. C. (2016). A multi-modal parcellation of human cerebral cortex. Nature, 536(7615), 171–178. 10.1038/nature18933

Groen, I. I. A., Dekker, T. M., Knapen, T., & Silson, E. H. (2022). Visuospatial coding as ubiquitous scaffolding for human cognition. Trends in Cognitive Sciences, 26(1), 81–96. 10.1016/j.tics.2021.10.011

Harrison, S. A., & Tong, F. (2009). Decoding reveals the contents of visual working memory in early visual areas. Nature, 458(7238), 632–635. 10.1038/nature07832

Haxby, J. V., Gobbini, M. I., Furey, M. L., Ishai, A., Schouten, J. L., & Pietrini, P. (2001). Distributed and Overlapping Representations of Faces and Objects in Ventral Temporal Cortex. Science, 293(5539), 2425–2430. 10.1126/science.1063736

Hong, H., Yamins, D. L. K., Majaj, N. J., & DiCarlo, J. J. (2016). Explicit information for category-orthogonal object properties increases along the ventral stream. Nature Neuroscience, 19(4), Article 4. 10.1038/nn.4247

Horn, A. (2016). *HCP-MMP1.0 projected on MNI2009a GM (volumetric) in NIfTI format* (p. 3172117 Bytes) [Dataset]. figshare. 10.6084/M9.FIGSHARE.3501911

Ito, M., Tamura, H., Fujita, I., & Tanaka, K. (1995). Size and position invariance of neuronal responses in monkey inferotemporal cortex. Journal of Neurophysiology, 73(1), 218–226. 10.1152/jn.1995.73.1.218

Kay, K. N., Weiner, K. S., & Grill-Spector, K. (2015). Attention Reduces Spatial Uncertainty in Human Ventral Temporal Cortex. Current Biology, 25(5), 595–600. 10.1016/j.cub.2014.12.050

Kay, K. N., Winawer, J., Mezer, A., & Wandell, B. A. (2013). Compressive spatial summation in human visual cortex. Journal of Neurophysiology, 110(2), 481–494. 10.1152/jn.00105.2013

Kleiner, M., Brainard, D., & Pelli, D. (2007). What’s new in Psychtoolbox*-*3*?*

Kravitz, D. J., Kriegeskorte, N., & Baker, C. I. (2010). High-Level Visual Object Representations Are Constrained by Position. Cerebral Cortex, 20(12), 2916–2925. 10.1093/cercor/bhq042

Kravitz, D. J., Vinson, L. D., & Baker, C. I. (2008). How position dependent is visual object recognition? Trends in Cognitive Sciences, 12(3), 114–122. 10.1016/j.tics.2007.12.006

Kriegeskorte, N., Mur, M., Ruff, D. A., Kiani, R., Bodurka, J., Esteky, H., Tanaka, K., & Bandettini, P. A. (2008). Matching Categorical Object Representations in Inferior Temporal Cortex of Man and Monkey. Neuron, 60(6), 1126–1141. 10.1016/j.neuron.2008.10.043

Lepsien, J., & Nobre, A. C. (2007). Attentional Modulation of Object Representations in Working Memory. Cerebral Cortex, 17(9), 2072–2083. 10.1093/cercor/bhl116

Niemeier, M. (2004). A Contralateral Preference in the Lateral Occipital Area: Sensory and Attentional Mechanisms. Cerebral Cortex, 15(3), 325–331. 10.1093/cercor/bhh134

Pelli, D. G. (1997). The VideoToolbox software for visual psychophysics: Transforming numbers into movies. Spatial Vision, 10(4), 437–442. 10.1163/156856897X00366

Pratte, M. S., & Tong, F. (2014). Spatial specificity of working memory representations in the early visual cortex. Journal of Vision, 14(3), 22–22. https://doi.org/10/gbf34r

Reithler, J., Peters, J. C., & Goebel, R. (2017). Characterizing object- and position-dependent response profiles to uni- and bilateral stimulus configurations in human higher visual cortex: A 7T fMRI study. NeuroImage, 152, 551–562. 10.1016/j.neuroimage.2017.03.038

Robinson, A. K., Grootswagers, T., Shatek, S. M., Behrmann, M., & Carlson, T. A. (2025). Dynamics of visual object coding within and across the hemispheres: Objects in the periphery. Science Advances.

Schwarzlose, R. F., Swisher, J. D., Dang, S., & Kanwisher, N. (2008). The distribution of category and location information across object-selective regions in human visual cortex. Proceedings of the National Academy of Sciences, 105(11), 4447–4452. 10.1073/pnas.0800431105

Serences, J. T., Ester, E. F., Vogel, E. K., & Awh, E. (2009). Stimulus-Specific Delay Activity in Human Primary Visual Cortex. Psychological Science, 20(2), 207–214. 10.1111/j.1467-9280.2009.02276.x

Silson, E. H., Groen, I. I. A., & Baker, C. I. (2022). Direct comparison of contralateral bias and face/scene selectivity in human occipitotemporal cortex. Brain Structure and Function, 227(4), 1405–1421. 10.1007/s00429-021-02411-8

Teng, C., & Postle, B. R. (2021). Spatial specificity of feature-based interaction between working memory and visual processing. Journal of Experimental Psychology: Human Perception and Performance, 47(4), 495–507. 10.1037/xhp0000899

Tong, F., Harrison, S. A., Dewey, J. A., & Kamitani, Y. (2012). Relationship between BOLD amplitude and pattern classification of orientation-selective activity in the human visual cortex. NeuroImage, 63(3), 1212–1222. 10.1016/j.neuroimage.2012.08.005

Vallat, R. (2018). Pingouin: Statistics in Python. Journal of Open Source Software, 3(31), 1026. 10.21105/joss.01026

Vogel, E. K., & Machizawa, M. G. (2004). Neural activity predicts individual differences in visual working memory capacity. Nature, 428(6984), 748–751. 10.1038/nature02447

Wandell, B. A., Dumoulin, S. O., & Brewer, A. A. (2007). Visual Field Maps in Human Cortex. Neuron, 56(2), 366–383. 10.1016/j.neuron.2007.10.012

Wang, L., Mruczek, R. E. B., Arcaro, M. J., & Kastner, S. (2015). Probabilistic Maps of Visual Topography in Human Cortex. Cerebral Cortex, 25(10), 3911–3931. 10.1093/cercor/bhu277

Wang, S., Rajsic, J., & Woodman, G. F. (2019). The Contralateral Delay Activity Tracks the Sequential Loading of Objects into Visual Working Memory, Unlike Lateralized Alpha Oscillations. Journal of Cognitive Neuroscience, 31(11), 1689–1698. 10.1162/jocn_a_01446

Watson, A. B., & Pelli, D. G. (1983). Quest: A Bayesian adaptive psychometric method. Perception & Psychophysics, 33(2), 113–120. 10.3758/BF03202828

Wolff, M. J., Jochim, J., Akyürek, E. G., Buschman, T. J., & Stokes, M. G. (2020). Drifting codes within a stable coding scheme for working memory. PLOS Biology, 18(3), e3000625. 10.1371/journal.pbio.3000625

Xu, Y. (2023). Parietal-driven visual working memory representation in occipito-temporal cortex. Current Biology, 33(20), 4516–4523.e5. 10.1016/j.cub.2023.08.080

Xu, Y., & Chun, M. M. (2006). Dissociable neural mechanisms supporting visual short-term memory for objects. Nature, 440(7080), Article 7080. 10.1038/nature04262

Yu, Q., & Shim, W. M. (2017). Occipital, parietal, and frontal cortices selectively maintain task-relevant features of multi-feature objects in visual working memory. NeuroImage, 157, 97–107. 10.1016/j.neuroimage.2017.05.055

Zhao, Y.-J., Kay, K. N., Tian, Y., & Ku, Y. (2021). Sensory Recruitment Revisited: Ipsilateral V1 Involved in Visual Working Memory. *Cerebral Cortex*, bhab300. 10.1093/cercor/bhab300

